# Evaluative Foraging: From Value to Action

**DOI:** 10.1101/2025.05.04.652128

**Authors:** Ana Clemente, Olivier Penacchio

## Abstract

**Short abstract:** Assigning hedonic value shapes behaviour across organisms, yet how these values translate into motivated behaviour remains unclear. We introduce *evaluative foraging* as a mechanism by which hedonic evaluation guides behaviour via iterative reward maximisation. This framework integrates hedonic evaluation with active inference, whereby expected free energy encodes wanting – the motivational drive toward rewarding outcomes – while variational free energy corresponds to liking – the hedonic impact of the outcome. Evaluative foraging unifies the mechanisms underlying behaviours ranging from allostasis to art appreciation, casting cognition and motivated behaviour as purposive engagement with the environment and its affordances.

**Long abstract:** Hedonic evaluation shapes behaviour. Yet, the mechanisms that transform hedonic values into motivated behaviour remain poorly understood, and key questions remain unanswered. How do habits, wanting and liking interact to generate behaviour? Do common principles govern behaviour across species, cognitive systems and behavioural complexity? We propose a unifying framework for understanding motivated behaviour that integrates hedonic evaluation with active inference, a theory in which perception, action and cognition emerge from the imperative to minimise free energy – a proxy for the discrepancy between actual and preferred states. We argue that expected free energy encodes wanting – the motivational drive toward rewarding outcomes – while variational free energy provides a measure of liking and disliking of resulting outcomes. This synthesis, which we term *evaluative foraging*, offers a principled explanation for how organisms navigate their environments based on a common mechanism of pleasure-driven learning and decision-making. It posits that motivated behaviour emerges from the iterative pursuit of maximising reward, with hedonic evaluation motivating and monitoring actions through wanting and liking mechanisms. We demonstrate how evaluative foraging captures a wide range of behaviours, from allostatic processes to sophisticated cultural endeavours such as art appreciation. Our integrative perspective enables testing the neurobiological implementation of the core constructs of active inference as the main components of hedonic evaluation, and fosters mutual enrichment between these fundamental approaches to behaviour. Ultimately, evaluative foraging reinforces the view that cognition is essentially about purposive engagement in adaptive interactions with environmental affordances and offers a unified framework for investigating cognition and behaviour.

## 1. Introduction

Motivated behaviour is usually examined through highly specialised lenses, focusing on idiosyncratic biological adaptations, neural processes and cognitive mechanisms. This leads to a proliferation of theories specific to particular fields or taxa, with limited prospects for unification. Yet, one of the main endeavours of cognitive psychology and neuroscience is to unveil general mechanisms of behaviour (Berridge, 2004; Friston, 2010), raising the fundamental question: Is there a common mechanism driving behaviour across the spectrum of complexity and species? For instance, is there a common underlying principle in a *C. elegans* orienting in a gradient of nutrients, an endotherm foraging to maintain metabolic balance and a human formulating complex plans to achieve distant goals?

When searching for unifying theories in biology and the behavioural sciences, evolution always emerges as a key principle (Gintis, 2007; Mesoudi et al., 2006). To be biologically meaningful, behaviour must be adaptive, enhancing the organism’s chances of survival and reproduction by improving its interaction with the environment. Behavioural mechanisms thus bear the imprint of selection across ontogenetic and phylogenetic levels (Duckworth, 2009; Snell-Rood, 2013).

Adaptive behaviour relies on evaluating the environment – its elements, relationships and affordances (Chiel & Beer, 1997; Schwarz, 2007). The fundamental neurobiological process of assessing stimuli as beneficial or harmful, rewarding or punishing, is referred to as *hedonic evaluation.* This process underpins the capacity to compare alternatives, make decisions and prioritise actions (Berridge & Kringelbach, 2015; Rosenthal, 2017). Whether selecting food, choosing a mate or engaging in leisure activities, organisms with a reward system are driven by the pursuit of rewards and avoidance of threats (Berridge & Kringelbach, 2008; Kringelbach, 2005; Rangel et al., 2008). This reward-seeking behaviour has been fundamental to brain evolution (Schultz, 2015); since the emergence of chordates (Cisek, 2022), neurons in the reward system compute value signals by integrating sensory input with interoceptive and executive information, reflecting the organism’s past experiences, states, goals and expectations (Rangel et al., 2008), thereby motivating and evaluating behaviour (Berridge & Kringelbach, 2013, 2015).

Motivated behaviour unfolds in three interrelated phases, each involving specific mechanisms and value signals (Berridge & Kringelbach, 2013; Pessiglione & Lebreton, 2015). In the appetitive phase, *wanting* mechanisms anticipate the reward of behavioural outcomes, guiding approach or avoidance, biasing perception towards salient features and modulating neural activity in perceptual and cognitive systems. During the consummatory phase, these motivational aspects integrate with executive and motor systems to implement behavioural choices. Finally, in the satiety phase, *liking* (and *disliking*) mechanisms evaluate the behavioural outcomes, signalling reward and informing future behaviour through learning (Dayan, 2022).^1^ Hedonic values are thus affective states that reward or punish behavioural decisions and inform future behaviour (Berridge & Kringelbach, 2015).

Yet, the mechanisms by which the information conveyed by hedonic values is transformed into behaviour are not fully understood. For example, how does the pleasurable taste of a fruit urge someone to eat it *and* someone else to cultivate an orchard? Whereas the former seeks immediate, simple reward, the latter engages in a long-term, risky and costly endeavour aiming for a future, complex reward. The picture gets even more complicated when multiple hedonic values conflict, as when animals explore new environments to seek novel and better resources despite facing a greater risk of predation. Hedonic evaluation explains the neuropsychological mechanisms of reward, but it does not explain how hedonic values are computed and translated into concrete actions. As a result, it fails to make specific predictions about future behaviour. This limitation arises because hedonic evaluation lacks a computational mechanism for selecting and implementing behavioural choices and, therefore, fails to establish causal relationships between hedonic value and subsequent behaviour.

Core adaptive mechanisms are rooted in the biological imperative to maintain homeostasis, driven by evaluative processes that compare the organism’s state with reference values (Cannon, 1929). This imperative provides a natural link to action, as deviations from homeostatic set points generate hedonic signals that motivate corrective behaviours to restore balance (Ramsay & Woods, 2014). For instance, organisms require precise glycaemia ranges, experiencing pleasure when approaching them (e.g. eating) and distress when deviating from them (e.g. experiencing hunger or indigestion). Importantly, the behaviour to approach such preferences depends on the organism’s states, needs and goals, as eating may be pleasurable when hungry but unpleasant when satiated (Berridge, 1991; Skov, 2023). Therefore, quantitative measures capturing the role of hedonic value in motivating behaviour, and thus a computational framework for motivated behaviour, must be grounded in comparisons with preferences and tailored to embodied and situated behaviour (Roth & Jornet, 2013; Shapiro, 2011).

While the imperative of maintaining homeostasis has long been recognised as central for living organisms, with the view that deviations trigger corrective mechanisms through reflexive actions (Cannon, 1929), recent approaches focus on the anticipatory nature of physiological regulation, a process called *allostasis* (Kleckner et al., 2017; Sterling, 2012). This operating principle is increasingly seen as the brain’s core function, with perception, cognition and action serving primarily to coordinate and regulate bodily needs (Theriault et al., 2025). Allostasis implies a crucial shift from a reactive to a *predictive* perspective on the brain, because a predictive brain anticipates future events, objects and demands (Clark, 2013). This has two key implications: First, the brain projects its preferred future outcomes, generating internal representations of the consequences of possible actions (Conant & Ross Ashby, 1970; Pezzulo & Cisek, 2016). Second, the brain commands anticipatory actions, which, according to its internal model, lead to its preferred outcomes (Wiener, 1948).

These two implications call for an *enactive* approach whereby the brain does not merely store internal models of the world to infer the causes of incoming information but, more fundamentally, develops models that enable a dynamic interaction between the organism and its environment. Such an interaction entails estimating conditions and affordances upon which the organism may *act* to fulfil its needs and goals. This perspective brings together two pivotal heritages in conceptualising and understanding perception, and the very *raison d’être* of perception itself (Friston, 2010): First, in Helmholtz’s view (von Helmholtz, 1866), perception is a process of inference of the causes of the incoming sensory input. The brain constantly combines top-down information from an internal generative model and bottom-up sensory evidence to estimate its environmental conditions. This view has been framed using a proper probabilistic setting in the *Bayesian brain hypothesis* (Doya et al., 2007; Gregory, 1980).

Second, in Gibson’s (Gibson, 1986) ecological theory of perception, the brain’s primary function is to perceive affordances, i.e. possibilities for action in the environment. In this action-oriented perspective, the brain is directly tuned to what the environment can offer to the organism without requiring complex internal processing.

To meaningfully explain and predict motivated behaviour, a mechanistic framework should therefore integrate predictive and enactive perspectives on the brain, proposing a probabilistic brain inference account of affordances in Bayes-optimal behaviour (Friston et al., 2006; Kording & Wolpert, 2006). Moreover, it needs to adopt an evolutionary perspective grounded in the simple axiom that an organism’s behaviour serves the imperative to survive in its environment (Friston, 2010). This axiom implies that organisms stay alive by adopting adaptive behaviours that bring them toward preferred states reflecting optimal living conditions within their ecological niche (Friston et al., 2012; Pezzulo et al., 2015). Friston’s principled approach to behaviour, *active inference*, provides a mathematical framework to operationalise such a process (Friston, 2010). Specifically, in active inference, behaviour reflects the imperative to *minimise free energy*, a proxy for the divergence between actual and preferred states.

In this article, we propose a biologically plausible computational framework for investigating behaviour that demonstrates how a broad range of scenarios can be accounted for by integrating hedonic evaluation with active inference. We show the potential of the proposed unified framework by modelling behaviour along a continuum, from basic allostatic processes shared by all organisms to more complex activities typically deemed uniquely human, such as art appreciation. This proposal allows us to bridge the gap and clarify the causal relationship between fundamental physiological processes involved in hedonic evaluation and action selection, offering a cohesive explanation for the motivational underpinnings of behaviour across a range of complexities. By framing hedonic evaluation within active inference, we propose a common ground for motivated behaviour that explains how organisms, from simple life forms to humans, navigate their environments, make decisions and pursue goals balancing immediate rewards with long-term survival and well-being.

We start by suggesting correspondences between the main components of hedonic evaluation, namely hedonic impact (core *liking* or *disliking*) and incentive salience (core *wanting*) (Berridge & Robinson, 1998), and the three components that guide policy choice in active inference, namely *variational free energy*, *expected free energy* and *habit* (K. Friston et al., 2017). From this correspondence, we propose a mechanism by which hedonic evaluation drives behaviour: *evaluative foraging*. Following this exposition, we examine why a continuum of behaviours should be expected across complexity and taxa, and illustrate evaluative foraging through selected scenarios to demonstrate its suitability for accounting for such a continuum. We argue that evaluative foraging ascribes complex cognitive functions such as appreciating art to the set of behaviours already accounted for by active inference, therefore welding core biological mechanisms and sophisticated cultural pursuits. This leads us to discussing how our framework can be empirically tested, providing a concrete basis for evaluating its validity. Finally, we discuss and contextualise evaluative foraging in relation to existing proposals and elaborate on the implications and contributions of the framework. To conclude, we emphasise the consequences and potential of a unifying, enactive and embodied computational mechanistic framework for understanding cognition and behaviour.

## 2. Evaluative foraging: Hedonic evaluation as active inference

As previously outlined, how hedonic evaluation, a core motivational process, translates into a framework for understanding and predicting behaviour remains undefined. There is no clear mechanism for computing distinct hedonic values, integrating them and guiding behaviour over time. To fill this fundamental gap and understand motivated behaviour, we propose mapping hedonic evaluation onto active inference, a principled theory that explains and predicts embodied cognition and behaviour (Pezzulo et al., 2024). The central move is to express the role of hedonic evaluation in monitoring and motivating free-energy minimisation.

Free-energy minimisation is at the core of active inference. The theory stems from the truism that organisms must maintain their bodily parameters within some limits to stay alive. This pursuit is framed using the concept of *surprise*^2^ (Pezzulo et al., 2024). Surprising observations (whether exteroceptive, interoceptive or proprioceptive) are those that deviate from reference observations, which are called *preferences* and reflect the range of values characteristic and beneficial for the organism (Friston, 2010). In active inference, preferences are modelled as *expectations* about observations reflecting states of the world aligned with the organism’s needs and goals. Surprise thus corresponds to *prediction errors* when such expectations are not met. Active inference posits that a single, unified principle drives behaviour: the imperative of minimising surprise, or, more precisely, minimising a tractable proxy for surprise called *free energy* (Pezzulo et al., 2024) (**Box 1**).

An organism minimises free energy through an inference process based on a *generative model*, namely a probabilistic mapping between hidden states of the world and their associated observations. The (true) states of the world are called *hidden* because the organism may not have direct access to the true unfolding process (the *generative process*) occurring in the world (**Box 1**). Instead, the organism relies on observations to infer the underlying states. Following the tradition of perception as inference – originating with Helmholtz and further developed in the *Bayesian brain* hypothesis (Doya et al., 2007; von Helmholtz, 1866) –, the inference process is conceived as an inversion of the generative model, estimating the most likely states given observations.

Active inference extends Bayesian theories by incorporating *action* into the inference process. The brain does not merely infer causes passively; it also generates actions to minimise discrepancies between predicted and actual observations. To minimise surprise and maintain preferred states, organisms continuously invert their generative model, comparing predicted with actual observations, to actively reduce the gap between them. This surprise-minimisation process drives both perception – updating beliefs as in perception as inference (von Helmholtz, 1866) – and action – modifying the environment to achieve preferred outcomes. Consequently, active inference is inherently enactive, integrating action into the inference process to enable adaptive behaviour in a dynamic world. For example, organisms cannot survive outside specific glycaemic levels, and deviations with respect to their preferred (physiologically optimal) range constitute surprising states. They cannot directly access their actual glucose index as reported by a physical glucometer but sense its effects (e.g. hunger) through interoceptive mechanisms involving the hypothalamus (Marty et al., 2007). Feeding is the action directed at restoring preferred glycaemic levels.

While active inference proposes a computational mechanism by which behaviour emerges from the drive to reach preferred states, hedonic evaluation provides organisms with information about the discrepancies between actual and desired states by assigning hedonic values to expected (wanting) and resulting (liking) behavioural outcomes to guide future actions (Dayan, 2022). Both the will and action of approaching preferences are positively signalled (e.g. the desire and pleasure of eating), whereas staying away or departing from preferences is negatively signalled (e.g. the displeasure of being hungry). In this proposal, we integrate hedonic evaluation and active inference to derive a general theory of how hedonic value motivates behaviour, and illustrate how the proposed framework, *evaluative foraging*, enables predicting behaviour. In the remainder of this section, we establish correspondences between the main components of hedonic evaluation and active inference, which form the foundation of our proposal, ultimately leading to evaluative foraging as a unified framework for understanding the core principles of motivated behaviour.

**Box 1.**
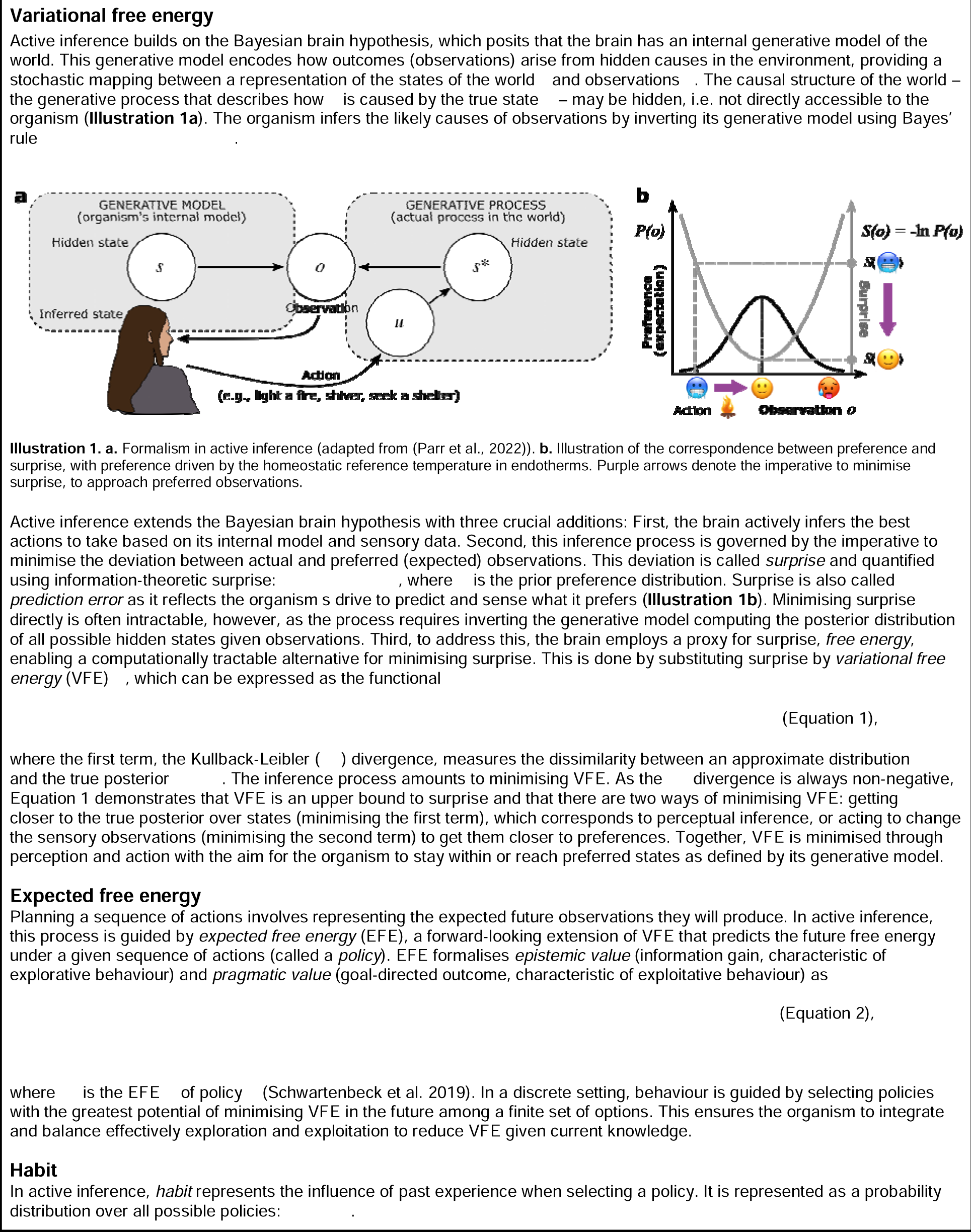
Formalisation of the active inference components of evaluative foraging.

### 2.1. Hedonic impact (core liking) as variational free energy

From the perspective of active inference, organisms seek states that best match their preferences (Friston, 2010). When an organism gets closer to its preferred (expected) states, it experiences less surprise and hence reduced *variational free energy* (henceforth VFE). From the perspective of hedonic evaluation, these states are rewarding (Berridge & Kringelbach, 2013; Pessiglione & Lebreton, 2015).

We propose that VFE can be considered as a quantitative measure of hedonic impact or core *liking* (**Box 1**). This hypothesised correspondence constitutes our proposal’s first and central component to build a framework that casts hedonic evaluation onto active inference.

As an illustration, consider a predator that expects to be in the state of satiation. When the predator is hungry, this discrepancy leads to prediction errors in its brain and is associated with high free energy. Through eating, this mismatch decreases, reducing VFE and thus eliciting a rewarding sensation. Similarly, a person listening to Western classical music may experience pleasure when a perfect cadence resolves to the tonic. This fulfilment of expectations following uncertainty corresponds to a transition to lower VFE which manifests as a pleasurable experience (Koelsch et al., 2019). In both examples, we propose that VFE quantifies the value of the outcome of the chosen behaviour and is neurobiologically implemented through hedonic impact (liking or disliking) mechanisms. Such a quantitative assessment is crucial to compare and motivate behaviour.

### 2.2. Incentive salience (core wanting) as expected free energy

While a reference measure for hedonic impact is necessary to understand motivated behaviour, it is not sufficient to predict it and thus explain how hedonic evaluation motivates behaviour. The reason is simple: VFE is retrospective; it evaluates incoming observations *a posteriori* and thus refers to past and present states. However, for planning and decision-making, an organism needs to infer the most effective future action or sequence of actions (hereafter referred to as *policy*) that generate observations yielding the most positive hedonic value signalling the lowest VFE. To infer optimal policies, the organism must predict sequences of future states and observations and, for each possible policy, estimate its resulting VFE, that is, compute its *expected free energy* (hereafter EFE) (**Box 1**). The prospective nature of EFE enables organisms to evaluate and select policies based on their impact on future states. EFE extends VFE to account for exploration, curiosity, planning and decision-making, making it central to active inference as a theory of goal-directed behaviour (Friston et al., 2015).

We propose EFE as a quantitative measure of incentive salience or core *wanting*, the second component of our proposal (**Box 1**). EFE consists of two key elements: *pragmatic value* and *epistemic value* (**Box 1**, Equation 2), formalising exploitation–exploration, a dual concept ubiquitous in biology across tasks and species (Hills et al., 2015).

Minimising the pragmatic term of EFE (Equation 2 in **Box 1**) maximises the pragmatic value, the expected utility or reward associated with a future state consequence of the organism’s policy. This aspect of EFE evaluates future outcomes (observations in the future resulting from policies) according to the extent to which they would fulfil the organism’s goal, namely how close they would bring the organism to its preference. By maximising pragmatic value, organisms avoid adverse outcomes and maximise immediate reward, therefore driving *exploitative* behaviour (K. Friston et al., 2017). The anticipation of eating a prey enhances the likelihood of hunting it for a predator. Similarly, the anticipation of enjoying live music motivates people to attend concerts. In other words, hedonic evaluation tells us that organisms want what they believe they will like, and active inference reveals the causal relationship underlying their choice of the policy expected to best achieve their desired outcome.

Minimising the epistemic term of EFE (Equation 2 in **Box 1**) maximises epistemic value, the expected information gain of each policy. This drives *explorative* behaviour by favouring policies that maximise knowledge about the world, resolving uncertainties and minimising prediction error. In our proposal, the epistemic component relates to incentive salience by driving the pursuit of information even in the absence of immediate reward (Schwartenbeck et al., 2019). A predator can explore new territories to improve access to higher-quality or more abundant prey, just as humans browse new music to enhance future musical experiences. This component accounts for the curiosity drive and the intrinsic pleasure of learning, extensively examined in the literature on flow, curiosity, motivation and related fields (Gottlieb & Oudeyer, 2018).

In summary, minimising EFE formalises incentive motivation by increasing the likelihood of choosing policies leading to outcomes with more positive hedonic value in the short and long terms. This minimisation can be naturally cast onto core *wanting* (Berridge & Dayan, 2021; Berridge & Kringelbach, 2015), capturing the dual motivational pull of securing rewarding outcomes and fostering learning. This dual pursuit is not mutually exclusive in the formalisation; pragmatic and epistemic values can be separately weighted and balanced, although the environmental affordances usually impose prioritisation constraints and a resulting trade-off between exploitation and exploration (Hills et al., 2015). The predator above might tend to explore more under pressure (when usual prey is scarce), and a composer might feel the need for further exploration of sonic materials to produce more satisfactory pieces.

### 2.3. Learned hedonic values as habits

In active inference, learning occurs at multiple time scales (Friston et al., 2016). Capturing long-term associations between expectations and outcomes (Seger & Spiering, 2011), *habits* emerge from a history of repeated behaviour, often resulting from consistently selecting policies that were beneficial in the past (Thorndike, 1911). As such, habit constitutes an essential behavioural drive, completing the interrelations between hedonic values and the constructs of active inference (**Box 1**). While EFE drives goal-directed behaviour, characterised by adaptive flexibility to environmental contingencies (Dayan, 2009), habit accounts for recurrent behaviour defined by past reinforcements even if unrelated to current hedonic values, being thus automatic, computationally efficient and inflexible (Dolan & Dayan, 2013).

Habit formation provides adaptive advantages, as habits may maximise efficiency in policy selection for recurrent scenarios, minimising energetic costs and risks when the available options are invariable and no further exploration is required or desired (Dolan & Dayan, 2013). For example, visiting a particular location at a particular time or season when prey is abundant can be greatly beneficial for a predator, and concert series subscriptions can facilitate regular music enjoyment for people, in both cases minimising energetic costs and risks to find them.

### 2.4. Motivated behaviour as evaluative foraging

The central idea of this proposal is that motivated behaviour can be mechanistically characterised as an interaction between habits, EFE minimisation (wanting) and VFE minimisation (liking). First, policy selection depends on the combination of habits (*E*, reflecting a history of policy selection and resulting hedonic impact: liking or disliking) and EFE (*G*, corresponding to wanting for available policies). Then, the selected policy leads to new observations, whose VFE (*F*, corresponding to liking or disliking the resulting outcomes) evaluates the effectiveness of the policy. In this way, habit and wanting define the prior expectations over policies, while their combination with liking determines the posterior for the selected policy, which in turn informs subsequent priors. We refer to **Box 2** for a detailed formalisation of evaluative foraging.

Evaluative foraging can thus be understood as navigating a free energy landscape (**Figure 1**), with action selection guided by habits and EFE as a compass, and with continual adjustments informed by the VFE resulting from each policy. At each step, the organism chooses a policy that optimises, with some stochastic variability, a balance between habit and wanting. After each policy, the organism evaluates the VFE of the resulting observations in form of liking or disliking. This hedonic evaluation then updates the active inference process, modulating habits and wanting while more generally refining the generative model that guides the organism’s actions. Behaviour thus unfolds iteratively, with each iteration consisting of these two interdependent stages (**Figure 1**).

**Figure 1.**
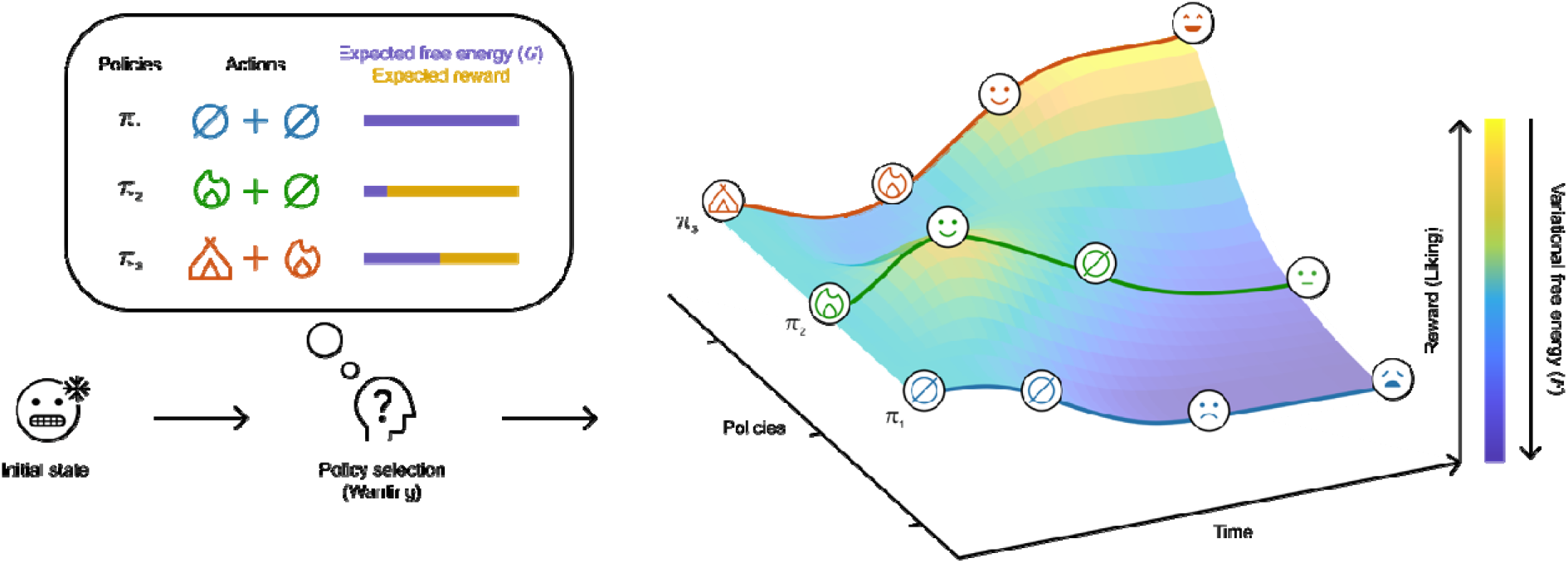
Evaluative foraging schematic: Free energy and reward landscape in thermoregulation. As endotherm organisms, humans must keep their body temperature within a specific range. A prolonged exposure to cold is life-threatening. The unpleasant sensation of cold increases the motivation (wanting) to restore body temperature. Available policies differ in their effectiveness, effort and time required to achieve this goal. For example, a hiker confronted with a cold sensation but not taking action (π_1_) would likely remain cold, further intensifying the unpleasant sensation (as VFE increases). However, she may act to reach a more beneficial state. For instance, she could light a fire (π_2_) to raise her body temperature quickly, providing an immediate reward (as VFE decreases), which is transient because her body temperature drops again when the fire extinguishes, once more eliciting the unpleasant sensation of cold (as VFE increases). Alternatively, she may consider the more elaborated, time-consuming and energy-expensive deeper policy (π_3_) of building a shelter (initially increasing VFE and hence discomfort) before lighting a fire, which ultimately leads to a greater and more lasting reward (as VFE decreases). Overall, the first policy (π_1_) would be disliked while the second (π_2_) would be mildly liked and the third (π_3_) would be strongly liked. By monitoring policies and their expected and experienced hedonic values at each time step, organisms can learn embodied associations between actions and outcomes, using this knowledge to guide future behavioural decisions.

Evaluative foraging casts behaviour as a balance between habits and EFE to minimise VFE, leading to various scenarios. In simpler cases, one component may dominate when the others are indifferent to the outcome of a new policy: e.g. habit may drive behaviour when all options are equally appealing and there is no incentive salience for learning – the distribution of EFE s flat. In more complex cases, a chosen policy may reflect a trade-off between competing components: e.g. when habitual behaviour conflicts with the drive for epistemic exploration. In the next section, we illustrate the scope of our proposal across species and behavioural complexities.

**Box 2.**
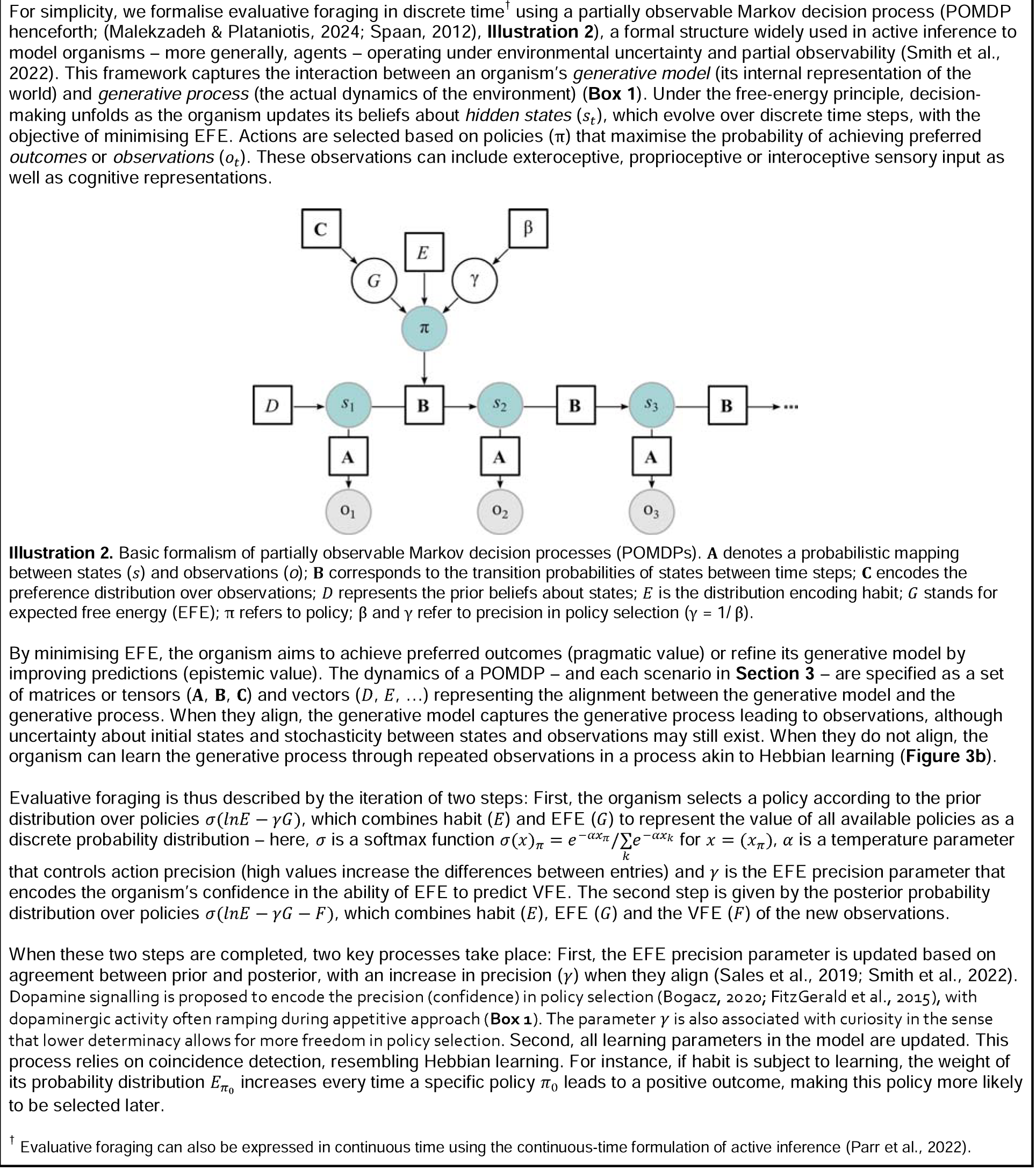
Evaluative foraging in discrete time using POMDPs.

## 3. A continuum of behaviours through evaluative foraging

In this section, we show how a continuum of behaviours can be captured within the unifying framework of evaluative foraging. We begin by examining why such a continuum should be expected across levels of behavioural complexity and biological taxa. Then, we demonstrate the scope of our proposal by simulating behaviour across four illustrative scenarios, highlighting its capacity to account for this continuum.

### 3.1. Rationale and evidence for a continuum

Evaluative foraging integrates hedonic evaluation into a single, unified model of action and perception (active inference) to describe a comprehensive range of behaviours that transcends biological taxa. But why should we expect a continuum in the first place? This enquiry is central to many disciplines, as it carries fundamental scientific and philosophical implications for understanding cognition and human nature, questioning whether there is a computational or neurophysiological mechanism of behaviour unique to *Homo sapiens*. If such a special mechanism exists, what does the break in the continuum look like, mean and entail for cognition? If a common mechanism drives a continuum of behaviours across scenarios and taxa, even sophisticated human pursuits and experiences traditionally deemed distinctly human (e.g. the so-called *aesthetic* experiences) would be grounded in the same principles as allostatic processes in simpler organisms such as worms. We can refine the question to resolve this conundrum as follows: Should we expect *wanting* and *liking* to promote and evaluate motivated behaviour, regardless of whether it be eating nutritionally valuable food (a primary reward satisfying biological needs) or consciously appreciating an artwork (a secondary reward without direct biological relevance)?

Scholars in empirical aesthetics and related disciplines have traditionally insisted on a negative answer to this question, based on the widespread but controversial belief that art appreciation is disconnected from other psychological and neurological processes (critically reviewed in (Nadal & Skov, 2024)). However, a central argument for a broadly shared basis of hedonic evaluation in motivated behaviour rests on substantial evidence that a wide array of pleasurable experiences (e.g. chocolate consumption, sexual activity, music listening) engages the same neural systems in humans, and that these systems are evolutionarily conserved and widely shared with other animals (Berridge & Kringelbach, 2013, 2015). This supports the hypothesis of a *common currency* for hedonic evaluation (Levy & Glimcher, 2012; Montague & King-Casas, 2007), whereby first and secondary rewards are processed within similar neural systems across species (Bartra et al., 2013; Sescousse et al., 2013).

From an evolutionary perspective, the function of *liking* is not to generate positive introspective experiences but to enable organisms to identify and respond to adaptive opportunities and threats (Pessiglione & Lebreton, 2015). Darwin (Darwin, 1872) anticipated this idea, proposing that emotional reactions are primarily selected for their adaptive value. This view has been substantiated by modern neuroscience, with extensive evidence for affective processing being underpinned by a core set of mechanisms shared across phylogeny (Anderson & Adolphs, 2014; LeDoux, 2012). Over time, cognitive and cultural pleasures have co-opted the hedonic circuitry initially evolved for sensory pleasures (Berridge & Dayan, 2021; Nguyen et al., 2021). Core hedonic values (affective signals) appeared first, followed by subjective and conscious evaluations computed by more complex hierarchical structures (Berridge & Kringelbach, 2015; Panksepp, 1998). Wanting might have evolved initially to pursue innate rewards, such as nutrients and mates, while liking likely emerged later to evaluate behaviour and inform subsequent wanting (Berridge & Dayan, 2021; Winkielman & Berridge, 2003). This would explain innate drives and exploration of novel environments, with learning resulting from the evaluation of such exploration. Accordingly, reward maximisation (EFE minimisation) would have been first initiated by uninformed predictions (hypothetical VFE) and would have been later informed by previous experiences (resulting VFE).

Crucially, we propose evaluative foraging as a common computational mechanism implemented through specific neurobiological processes. While the framework suggests that the principle of reward maximisation spans behaviours and species, the scenarios and brain structures underlying hedonic evaluation differ between taxa. More complex brains enable more refined cognitive abilities, hierarchical processing and sophisticated computations, such as deeper policy evaluation and planning; for example, *C. elegans*, whose central nervous system is made of 302 neurons, use simple exploration strategies for food search (Calhoun et al., 2014), while orcas plan and coordinate collective hunts (Pitman & Durban, 2012). Furthermore, distinct brain structures subserve similar functions in different species (Jarvis et al., 2005); for instance, executive functions involved in behavioural optimisation are associated with activity in the mammalian prefrontal cortex (Friedman & Robbins, 2022) and the avian nidopallium caudolaterale (Güntürkün & Bugnyar, 2016).

Notwithstanding specific differences, the neural machinery underpinning hedonic evaluation is conserved across phylogeny. Substantial converging evidence supports homology across major vertebrate lineages (O’Connell & Hofmann, 2011), and the emergence of the mesolimbic reward system predates the divergence of chordates (Yamamoto & Vernier, 2011). Biogenic amines (e.g. dopamine) are essential for reward processing across animal phyla (Barron et al., 2010), motivating behaviour in vertebrates and invertebrates (Costa & Schoenbaum, 2022; Unoki et al., 2006), and the insect reward system has a modular structure that broadly parallels the mammalian system in functional organisation, with distinct brain circuits for reward learning, wanting and liking (Perry & Barron, 2013).

Together, these functional, phylogenetic and neurophysiological similarities suggest a unified mechanism of motivated behaviour across scenarios and taxa. Hedonic evaluation mechanisms evolved to motivate and evaluate adaptive behaviour and are conserved across species (e.g. *C. elegans*, *Homo sapiens*), scenarios (e.g. eating fruit, cultivating an orchard) and stimuli (e.g. food, sex, artworks). Active inference explains this continuum from first principles, with behaviour driven by the imperative to maintain preferred states shaped by evolutionary adaptations to ecological niches. By integrating hedonic evaluation within active inference, evaluative foraging provides the computational mechanism through which reward maximisation drives and monitors behaviour in organisms with a reward system, situating the commonalities discussed above within a continuum of behaviours and taxa.

### 3.2. Instantiating the continuum through four scenarios

In this section, we illustrate evaluative foraging through four behavioural scenarios strategically chosen to demonstrate the relevance and scope of the framework. The models’ architecture and the computational mechanisms governing behaviour are the same across scenarios. This consistency substantiates evaluative foraging as accounting for a behavioural continuum. Only the model parameters are specific to each scenario.

Before turning to these scenarios, we outline the principal elements of active inference models. The following set of mappings defines the generative models: The first matrix or tensor (**A** in **Box 2**) maps hidden states to observable outcomes. This mapping is probabilistic, providing the likelihood of a particular observation given the true state of the world. For example (**Figure 1**, **Illustration 1b** in **Box 1**), a state of hypothermia leads to the sensation of cold most of the time. The second mapping (**B** in **Box 2**) describes the probabilistic transitions between hidden states from the present to the next time step, depending on each action taken. In our example, not moving keeps the agent in hypothermia, whereas looking for shelter entails a probability of transitioning to normothermia of 0.8. The preference matrix (**C** in **Box 2**) encodes the agent’s desired outcomes as probability distributions, shaping EFE and the VFE landscape. In this example, the hiker (adaptively) prefers staying in the reference (homeostatic) state of normothermia, which translates into positive hedonic values (wish, reward) for normothermia state and negative hedonic values (avoidance, punishment) for hypo- and hyperthermia.

In the POMDP-based scenarios below, behaviour emerges dynamically from the interaction between the components of evaluative foraging. Evaluative foraging is characterised as the stochastic evolution of the organism’s states and observations, driven by the combined influence of all POMDP factors (including observation mappings, state transitions, habits and preferences) constraining the organism’s choices in response to its environment. The code implementation and detailed description of each scenario are publicly accessible at https://osf.io/v39mb/?view_only=852e26bf70da4f7aa919ea41fb20fcd5.

#### 3.2.1. Scenario 1: Allostasis

Through simulations, we show how evaluative foraging accounts for allostasis in a simple setting: glucoregulation (Roh et al., 2016). An organism can be in one of three physiological states: hypoglycaemia, normoglycaemia or hyperglycaemia. It estimates these states through interoceptive observations: the sensations of hunger, satiety and indigestion, respectively. The mapping between hidden states and sensations can be noisy: a true state does not always align perfectly with its usual sensation (e.g. hypoglycaemia can be perceived as satiety). The organism iteratively chooses between two policies: eating increases the likelihood of transitioning to a higher glycaemic state (from hypoglycaemia to normoglycaemia and from normoglycaemia to hyperglycaemia) whereas resting increases the likelihood of transitioning to a lower glycaemic state (from hyperglycaemia to normoglycaemia and from normoglycaemia to hypoglycaemia). The organism strongly prefers satiety (normoglycaemia). Through evaluative foraging, the organism’s feeding cycle emerges from the imperative to reach its preferred state and avoid surprise (**Figure 2**).

**Figure 2.**
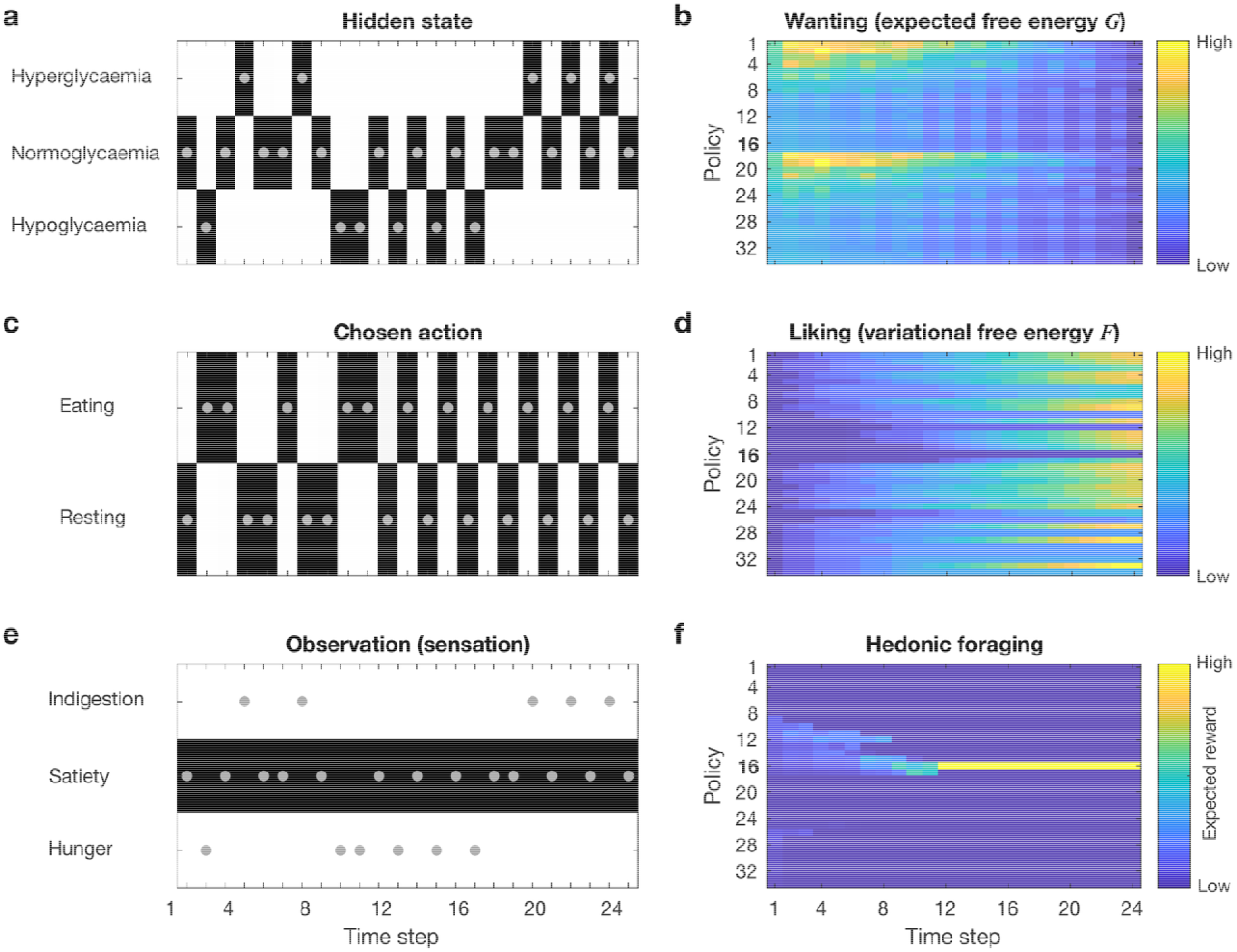
Allostasis (glucoregulation). In this instance of glycaemia regulation, the organism starts in normoglycaemia (*t*_1_). When its glycaemic level decays (*t*_2_), it enters a hypoglycaemic state and hence feels hungry, so it starts eating (*t*_3_) to reach a normoglycaemic state and feel satiated. Eating in normoglycaemia (*t*_4_) leads to hyperglycaemia, with a sensation of indigestion. Resting (*t*_5_) leads to progressive glycaemic decay until reaching the preferred normoglycaemic state, which progressively decays into a hypoglycaemic state, triggering eating again (*t*_6_). The left panels show how a feeding cycle emerges from the goal of maintaining normoglycaemia as long as possible. Dots represent the hidden state (**a**), the chosen action (**c**) and the resulting observation (**e**) at each time step. The process is not deterministic, so the organism can, for example, keep eating even when satiated. Policy choices are driven by evaluative foraging, with the greatest reward resulting from eating when feeling hungry. The right panels show the dynamics of wanting (**b**), liking (**d**) and evaluative foraging (**f**) – with lower EFE (*G*) corresponding to higher expected reward (wanting) and VFE (*F*) corresponding to actual reward (liking) –, where the choices converge into a stable policy (π_16_) yielding maximum global hedonic value.

#### 3.2.2. Scenario 2: Statistical learning and preference formation

Statistical learning is the process by which an organism implicitly learns regularities in sensory input, shaping future expectations (Turk-Browne, 2012). When an input aligns with learned patterns, it typically elicits a positive affective response because sensory–cognitive systems are suited to processing familiar features efficiently (Bornstein & Dagostino, 1992; Reber et al., 1998). Statistical learning thus subserves *mere exposure* effects, tendencies to develop liking for stimuli as a function of familiarity (Zajonc, 1968). This phenomenon is shared with other animals and influences the hedonic evaluation of a broad range of stimuli, from food (Birch & Marlin, 1982) to music (Pereira et al., 2011) and artworks (Reber et al., 2004).

This scenario exposes a link between statistical learning and preference formation. An organism freely chooses among three equally preferred locations: two contain stimulus A while the other contains a different but equally rewarding stimulus B. Without any incentive to favour a specific location, stimulus A is twice as likely to be encountered. Greater exposure reinforces preference for stimulus A over B, simply because its associated reward is better known. As the organism progressively learns the stimulus frequency and location, its pursuit of reward creates a feedforward loop whereby more encounters with the preferred stimulus strengthens the preference for it, fostering future encounters. This behaviour ultimately leads to establishing a habit of visiting the locations typically containing the preferred stimulus.

#### 3.2.3. Scenario 3: Bias and curiosity

Hedonic evaluation is context-dependent (Berridge, 1991). Through statistical learning, organisms identify regularities in spatial and temporal co-occurrences (Turk-Browne, 2012) and develop preferences for familiar patterns (**Scenario 2**). Consequently, preferences for specific stimuli can be *biased* by their co-occurrence with other stimuli or contextual features with enhanced evaluative salience (Lopez-Persem et al., 2016). For instance, the *white cube effect* refers to a tendency to prefer artworks displayed in museums over those encountered in public spaces (Gartus & Leder, 2014). However, *curiosity* (Gottlieb & Oudeyer, 2018; Kidd & Hayden, 2015) can counteract such biases. Evaluative foraging incorporates curiosity via the epistemic term of EFE, fostering exploration to reduce uncertainty (Schwartenbeck et al., 2019).

In this scenario, an organism (e.g. a museum visitor) can choose between two contexts leading to equally pleasurable stimuli (e.g. particular artworks). The organism prefers one of the contexts (e.g. museum over metro station) because it has statistically, culturally or socially learned that stimuli in this context are usually more rewarding than those in the other context. Incentive motivation (*G* in **Box 1**) or habit (*E* in **Box 1** and **Box 2**) to visit the preferred context more often bias the hedonic impact (*F* in **Box 1**) of stimuli in that context, enhancing the preference for that stimulus (**C** in **Box 2**) or the habit of visiting that context (*E* in **Box 1** and **Box 2**). **Figure 4** demonstrates that more curious organisms visit less preferred contexts more often (panel **4a** vs panel **4b**), which results in bias reduction as the stimuli can be evaluated in their own right.

**Figure 3.**
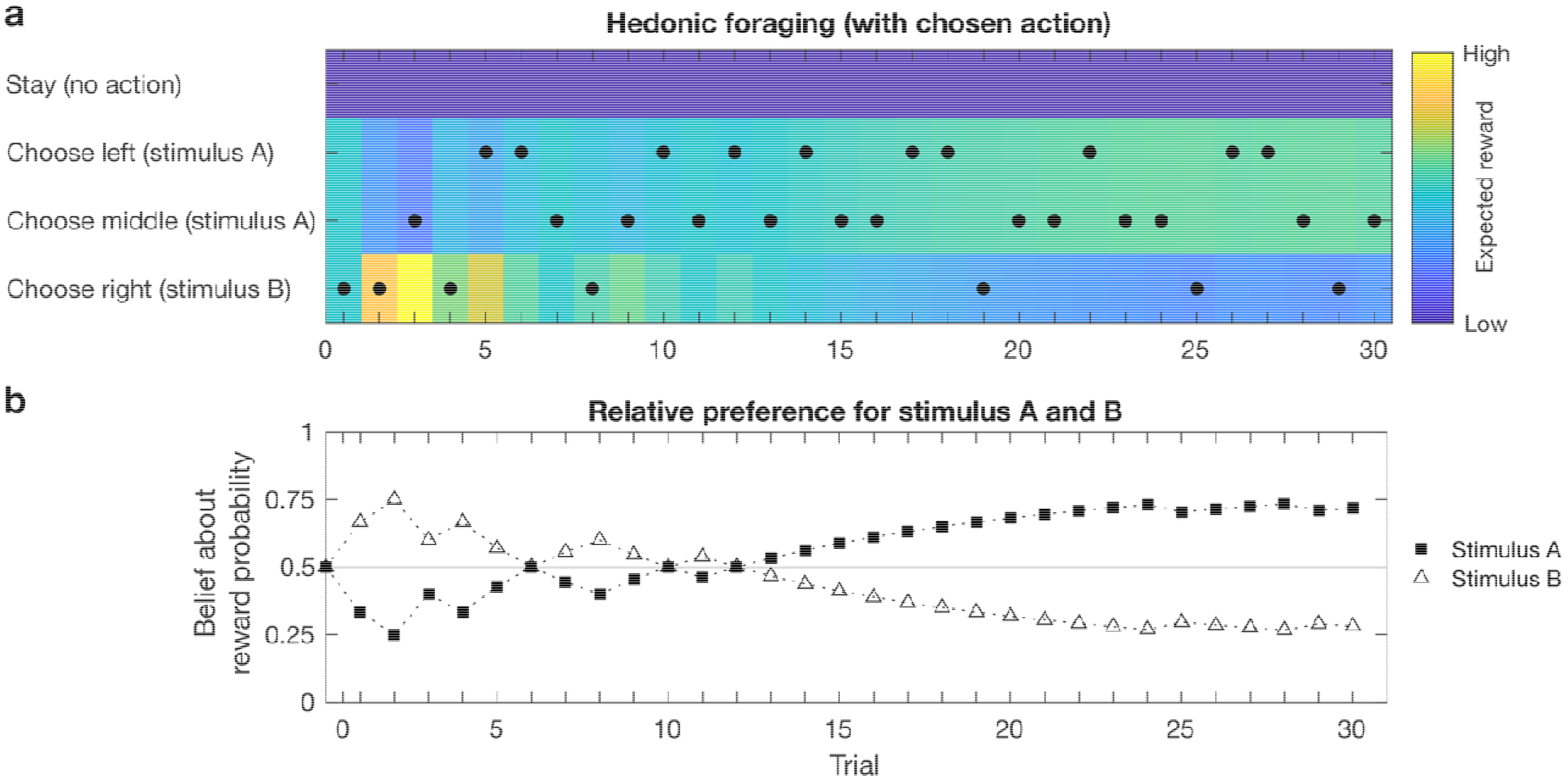
Statistical learning and formation of familiarity-driven preferences. In this example, the organism has no initial preference between stimulus A and B, leading to random choices. Over time, it learns stimulus frequency and location: A appears in 2/3 of locations while B does in 1/3. Despite initially sampling B (*t*_0_), the organism quickly infers that A is more frequent and shifts its preference accordingly, increasing visits to A-associated locations (*t*_>1_). Evaluative foraging drives behaviour by selecting policies (dots in **a**) that maximise reward according to updated beliefs about reward probability (trajectories for each stimulus in **b**).

**Figure 4.**
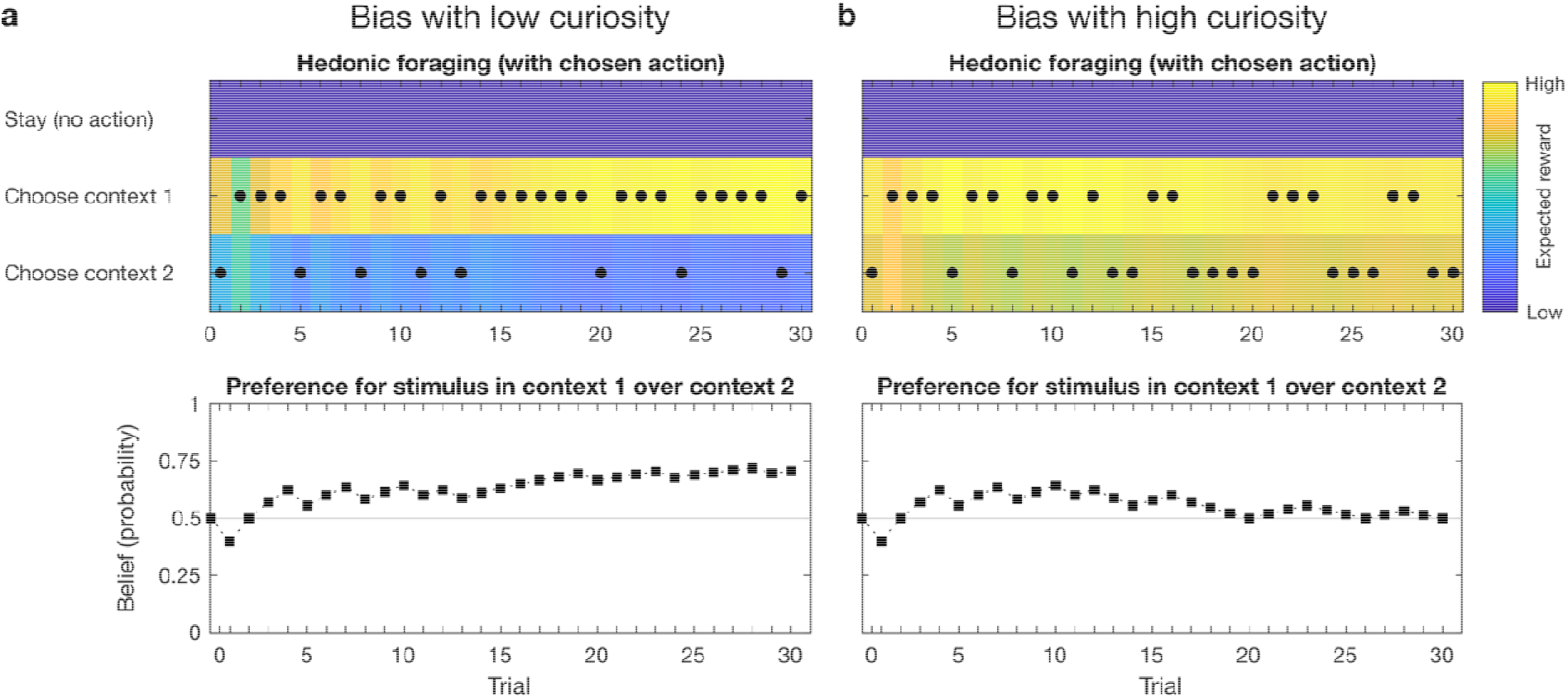
Interplay between bias and curiosity. In this example, contexts 1 and 2 contain the same stimuli. The organism has an initial bias (preference) for context 1, even if it starts visiting context 2 due to stochasticity in policy choice (*t*_0_). Evaluative foraging guides behaviour, with policy selection (top panels) maximising reward based on updated beliefs about reward probability (bottom panels). **a.** When curiosity is low, the organism visits context 1 more frequently (top panel) because it believes this context yields greater reward, in turn reinforcing this belief (bottom panel). **b.** High curiosity counteracts this bias, as exploration allows the organism to experience the same stimuli in both contexts and evaluate them in their own right. Increased exploration reveals that stimulus value is context-independent, leading to more balanced use of contexts (top panel) and reduction of prior biases (bottom panel).

#### 3.2.4. Scenario 4: Insight, *Erlebnis* and “Aha” experiences

Ambiguous stimuli elicit unstable percepts (Klink et al., 2012). From the perspective of predictive coding within active inference, uncertainty generates prediction error (Doya et al., 2007; Ross & Hansen, 2016). Organisms seek to resolve uncertainty and hence reduce prediction error, deriving reward from the process (FitzGerald et al., 2015). When understanding occurs suddenly, with a feeling of ease and certainty (Topolinski & Reber, 2010), usually requiring only a few observations and favoured by curious behaviour (K. J. Friston et al., 2017), it is called *insight*, *Erlebnis* or “Aha” experience. Gaining understanding or insight can occur exogenously (through external cues or explicit information) or endogenously (through internal reasoning or exploration), each with a distinct probability of success. Organisms vary in their preference for seeking insight exogenously or endogenously and in their ability to gain insight with or without external aid (see (Kounios & Beeman, 2014) for a review in humans and (Shettleworth, 2012) for a discussion in other animals).

This scenario shows how individual ability and relative preference for exogenous or endogenous insight guides behaviour. It can be situated, for example, in a museum where visitors choose whether to consult the labels accompanying the artworks (**Figure 5**).

**Figure 5.**
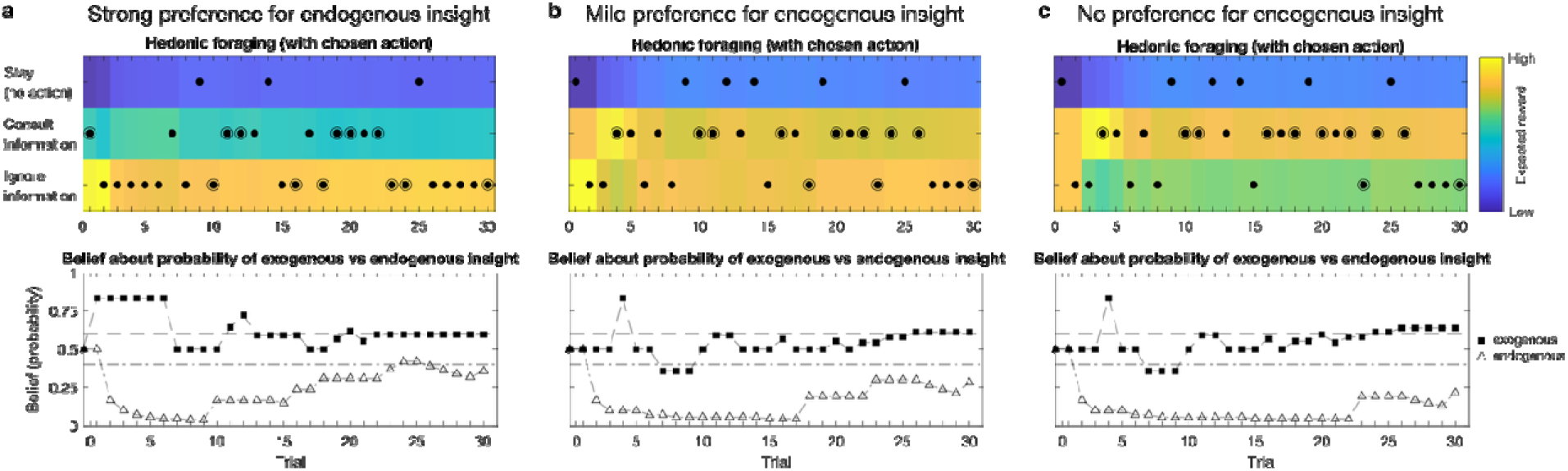
Preference for endogenous vs exogenous insight. An organism is presented with an ambiguous stimulus and acts to resolve its uncertainty. This can be achieved through two strategies: exogenously by consulting external information, which increases the probability of insight, or endogenously, relying on internal processing with a lower probability of insight. Organisms exhibit individual variability in their preference for endogenous over exogenous insight, and distinct ability (probability) to gain insight (circled dots). For simplicity, ability is constant: 0.6 for exogenous and 0.4 for endogenous insight. Evaluative foraging captures how such traits determine individual behaviour (dots represent chosen actions in the upper panels, while the lower panels depict the evolution of beliefs about the relative probabilities of gaining insight). **a.** When the preference for endogenous insight is stronger than the preference for exogenous insight, the expected reward outweighs the risk of not getting insight, leading the organism to disregard external information and trust its own skills. **b.** When the preference for endogenous insight does not compensate for the risk of not getting insight, both strategies are balanced. **c.** In absence of preference for endogenous insight, the organism prioritises gaining insight, relying more on exogenous facilitation.

## 4. Testing and applying evaluative foraging

The value of any theory or model resides in its heuristic utility and predictive power – its ability to explain existing phenomena and anticipate new data (Lipton, 2005). We have shown that a common underlying principle can account for a wide range of motivated behaviours, from a hungry nematode foraging in a mud pool to the execution of complex plans by humans. This breadth raises a standard concern for unifying theories: whether generality comes at the expense of testability.

In this section, we start by clarifying how a single overarching principle need not preclude an understanding of the diverse biological expressions of motivated behaviour. Then, we explain how computational parameters can be extracted from empirical data to compare the dynamics of the core components of active inference and hedonic evaluation, offering a data-driven test of the evaluative foraging framework. Together, these steps establish the framework as both empirically tractable and biologically interpretable.

### 4.1. A single unifying framework with specific predictions

How can evaluative foraging capture and predict goal-directed behaviour across a continuum of contextual and biological complexity while resting on a single unifying principle? Within this framework, the core constructs and operational principles of active inference remain constant, with only the modulating parameters – including the fine-grained form and complexity of the generative model – varying to suit each organism and context. This preserves mechanistic continuity while accounting for biological and contextual diversity. Differences in behavioural complexity are expressed through hierarchical generative models that encode internal and environmental structure and dynamics across temporal and conceptual scales. Successive hierarchical levels generate predictions about hidden causes, spanning raw sensations to extended objectives, thereby supporting behaviour spanning from reflex arcs to sophisticated, goal-directed actions.

Consider the examples of a predator confronting potential prey and a human listening to music. Despite their differences, these experiences share common dynamics of free-energy minimisation and reward maximisation, supported by similar neurophysiological systems (**Section 3**). Yet, predator–prey interactions and human music listening involve behavioural experiences that can be brief or prolonged and are strongly context dependent, with human experiences often characterised by especially rich and multifaceted emotional engagement.

Central to these experiences are a constellation of factors, including previous knowledge (e.g. whether the prey is palatable, or whether the music is familiar), memories (e.g. linked to important events or people) and the broader context, both exogenous (e.g. food scarcity) and endogenous, such as the current physiological (e.g. hunger) and psychological state (e.g. sadness). Purpose and expectations (e.g. mood regulation) as well as individual and social circumstances (e.g. shared with someone meaningful) further shape these experiences.

Together, these interacting factors – from broad considerations to fine-grained individual differences – determine the structure and parameters of the generative models. Such detailed and flexible mappings endow evaluative foraging with capacity to make specific predictions about factors shaping behaviour, therefore enabling direct examination of the theory and its utility to address relevant questions about the lives and behaviour of organisms and their neural bases. The remainder of this section focuses on explaining how this framework can be empirically tested and applied.

### 4.2. Fitting empirical data through computational phenotyping

A central concept for testing evaluative foraging is *computational phenotyping*. At the core of any parametric model of behaviour, lies a mapping between model parameters and observed behaviour. Computational phenotyping consists in inverting this mapping to recover parameters from empirical data (Schwartenbeck & Friston, 2016). This technique has served to characterise individual variability across domains, including social behaviour (Xiang et al., 2012), personality, developmental and clinical neuroscience (Patzelt et al., 2018), cognition (Schurr et al., 2024) and genetics (Drouin et al., 2016). It is particularly influential in applications of active inference, where recovered parameters retain direct cognitive interpretability (Schwartenbeck & Friston, 2016; Smith et al., 2021), offering double benefit: prediction of behaviour from empirically estimated parameters, and interpretation of these parameters in terms of core motivational and cognitive processes.

To illustrate, predators often prefer cryptic prey because conspicuous signals advertise defences such as toxicity (Mappes et al., 2005; Penacchio et al., 2024). Within active inference, this preference reflects minimisation of surprise, as sampling defended prey generates aversive outcomes (Penacchio et al., 2025). The strength of this preference is captured by the matrix (**Box 2**) and can be estimated experimentally, for example by manipulating prey bitterness (Skelhorn & Rowe, 2006). Such inference can be performed at the individual or group levels, for instance comparing juvenile and adult predators and testing hypotheses about innate versus learned avoidance of conspicuous prey. Avoidance behaviour is context-dependent: preference for cryptic prey diminishes when food is scarce, as the imperative to prevent hunger overrides the drive to avoid toxins (Barnett et al., 2007). This added complexity can be captured with a two-level hierarchical model in which the upper tier infers the current context and sets overarching goals, while the lower tier selects context-appropriate actions – with the upper-level matrix encoding the value of high-level outcomes (e.g. securing calories) and the lower-level matrix expressing immediate preferences (e.g. avoiding toxins). This approach yields concrete, testable predictions (e.g. adults will be more reluctant than juveniles to tolerate some toxins) and can illuminate sources of individual variability and species-specific strategies, as commonly examined in behavioural and cognitive ecology.

Music also provides a rich model domain for studying cognition and behaviour. **Scenario 4** illustrated how skills and preferences interplay to drive behaviour and how these can be acquired (learned) through reward maximisation, explaining the interaction between curiosity and expertise. Greater expertise can be characterised by a more deterministic mapping between states and observations and more deterministic transitions between states, statistically learned through extensive exposure and practice. For instance, individual sound events, their voice-leading and harmonic relationships in a Western classical musical piece are more precisely defined for classical musicians than for aficionados (Schlaug, 2015; Zatorre et al., 2007). In active inference, perceptual accuracy can be expressed as a higher probability of correspondence between actual sounds (states) and their percepts (observations) in the tensors of the generative model (**Box 2**). Predictive accuracy in experts can be expressed as higher probabilities for particular transitions between events (states) in tensor (**Box 2**). More refined and established preferences (**C** in **Box 2**) for how music should unfold explains why musicians experience greater surprise when harmonic or voice-leading relationships are violated, and why they are less tolerant to mistakes (Hernández et al., 2019). These mappings can be hierarchical, such that lower-level auditory features (e.g. pitch, timbre) are integrated into increasingly abstract representations (e.g. tonal structure, musical style), with each level constraining inferences at the one below.

To summarise, computational phenotyping within evaluative foraging enables precisely formulated, empirically testable questions linking organism, task and context. Model parameters inferred from behaviour can be interpreted in terms of curiosity, learning and valuation, grounding abstract constructs in measurable data.

### 4.3 Closing the loop: Active inference predictions and hedonic evaluation markers in iterative interplay

At the core of the evaluative foraging framework is the correspondence of EFE with wanting and of VFE with liking, and how their interplay drives behavioural trajectories. Testing the framework therefore requires examining whether the dynamics of computational quantities align with independent measures of hedonic values.

Computational phenotyping yields estimates of generative-model parameters and trial-by-trial trajectories of habits, EFE and VFE during behaviour. In parallel, liking and wanting can be assessed via neural activity (e.g. EEG, MEG, fMRI), autonomic physiology (e.g. heart rate, pupil dilation) and behavioural reports (Berridge & Kringelbach, 2015; Schultz, 1998). Evaluative foraging predicts iterative, causal interactions, whereby wanting guides policy selection and liking evaluates outcomes, feeding forward to shape future habits and preferences. The central test lies in comparing these two routes: do the dynamics of EFE and VFE inferred from behaviour covary with independently measured indices of wanting and liking?

**Figure 6** schematises this process using pictorial creation as an example. An artist iteratively observes, evaluates and acts upon the evolving work, with each action producing new sensory input that updates evaluation and subsequent choice. Computational phenotyping enables estimation of individual generative-model components (**A**, **B**, **C**, *D*, *E*) from which the dynamics of VFE and EFE are derived. These trajectories can then be compared with the dynamics of neural and behavioural markers of liking and wanting. Causal hypotheses can be further tested using experimental manipulation or formal causality analyses.

**Figure 6.**
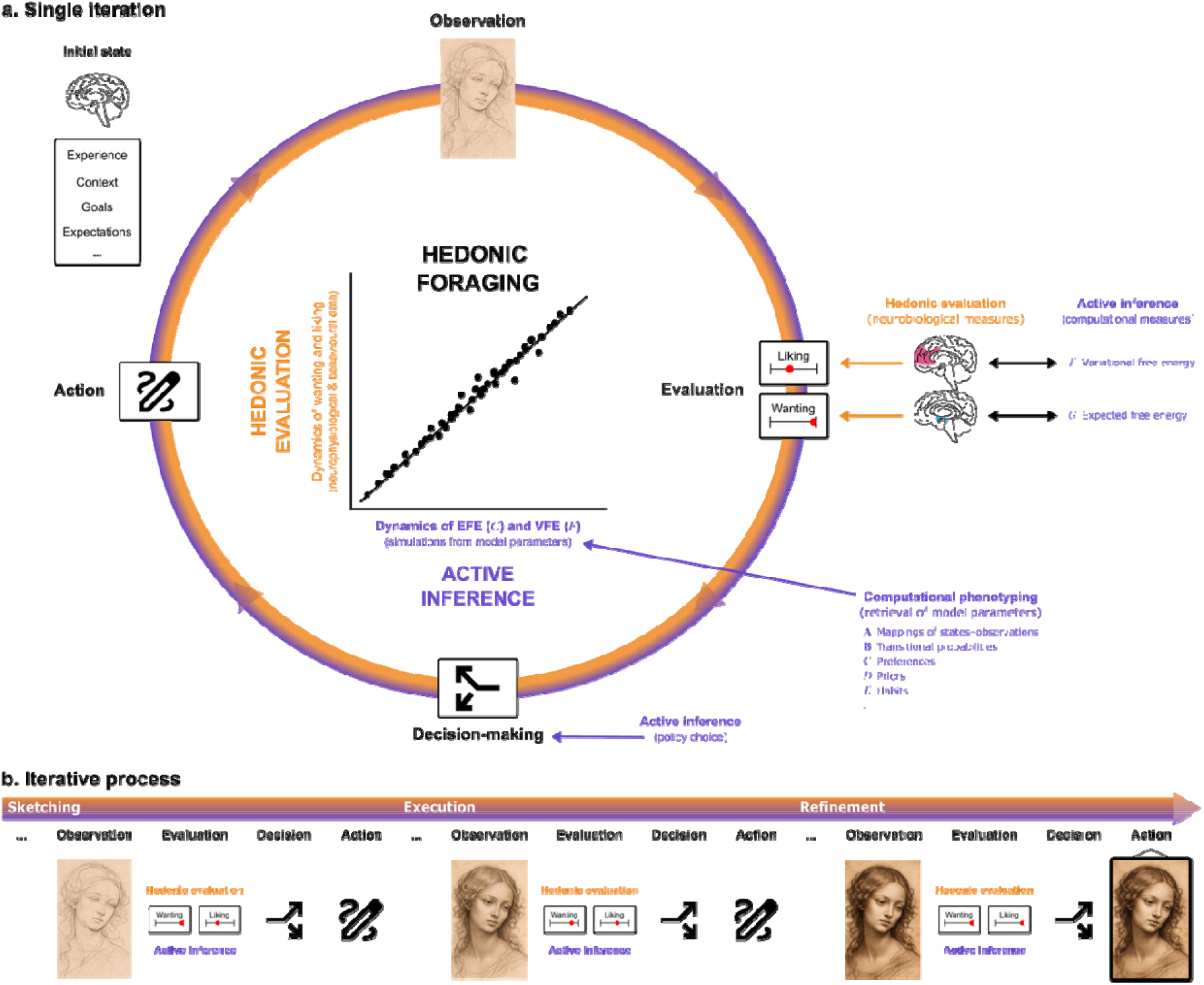
Testing evaluative foraging: Schematics of evaluative foraging as an iterative mechanism of monitoring and pursuit of free-energy minimisation as reward maximisation. In this example, an artist draws a Renaissance-style portrait. This process is hierarchically structured, spanning evaluations from single strokes to full gestures, motifs, sections, layers, composition and stylistic consistency. We show excerpts from three phases: *sketching* main lines to establish composition, proportions and placement; *execution*, during which forms, tones and textures are developed; and *refinement* of details to enhance contrast and balance. Panel a illustrates a single iteration, while panel b shows how these causal loops unfold over time. The iterative cycle starts by considering the initial state, including previous experience and current context, goals and expectations. Each iteration involves *observation* and *evaluation* of the current state of the creative process and product, followed by selection and execution of the next action. Within evaluative foraging, computational evaluation yields hedonic values that inform action selection: how closely current observations match preferred observations (liking, corresponding to VFE minimisation) and how much candidate policies are expected to bring future observations closer to those preferences (wanting, corresponding to EFE minimisation). Accordingly, the artist acts to produce a new observation, initiating a new iteration. This process terminates (e.g. the work is framed and displayed) when she deems the outcome satisfactory. Indices of brain activity (e.g. BOLD signal) most specifically associated with wanting (e.g. ventral tegmental area and nucleus accumbens; blue) and liking (e.g. prefrontal cortex and anterior cingulate cortex; pink) at each time step can be interpreted as implicit indicators of evaluative mechanisms (Berridge & Kringelbach, 2015; Schultz, 1998) and contrasted with explicit judgements (ratings; white boxes). Computational phenotyping enables retrieval of individual generative-model parameters (e.g. **A**, **B**, **C**, *D*, *E*, reflecting the situated internal and external states and operationalising factors such as expertise) that influence policy selection. From these parameters, the dynamics of VFE (*F*) and EFE (*G*) and are derived via simulations. These computational dynamics can then be compared to the neurobiological dynamics captured through neurophysiological and behavioural indices of liking and wanting. This enables a direct test of the alignment between wanting and EFE (*G*), and between liking and VFE (*F*), thereby probing the core predictions of evaluative foraging. Causal relationships between hedonic evaluations and behavioural decisions can be examined using formal causality analyses or experimental manipulations that target evaluative processes or the resulting observations.

To conclude, a single inferential mechanism grounded in free-energy minimisation can generate a wide range of behavioural patterns while accommodating contextual and biological diversity through parameterisation and hierarchy. The framework is empirically tractable: predicted trajectories can be directly contrasted with independent measures of hedonic evaluation, enabling a principled test of the correspondence between computational and neuropsychological constructs. By embedding hedonic evaluation within active inference, the framework links universal affective dynamics to individual- and species-specific behaviour.

Parameters such as precision, preference, learning rate and epistemic drive can be empirically estimated, yielding predictions relevant to motivation, learning and adaptation across domains – from allostasis to aesthetics. In the next section, we consider what such tests imply for empirical support of active inference more broadly, and contextualise our framework.

## 5. Contextualising evaluative foraging

In this section, we first review theoretical frameworks and computational models that explore the relationships between hedonic value – along with the related concepts of emotional valence and affective value – and motivated behaviour. We then contrast active-inference-based approaches with alternative paradigms, highlighting their principles and relevance to evaluative foraging. To conclude, we emphasise the implications of integrating hedonic evaluation and active inference for both frameworks and for the questions motivating this work.

### 5.1. Contrasting the evaluative foraging framework

#### 5.1.1. Emotion, affect and epistemic curiosity

Theories and models built on active inference aiming to establish principled links between hedonic value and behaviour have primarily focused on the subjective emotional impact of experiences; specifically, on *emotional valence,* the positive or negative quality of emotion. (Joffily & Coricelli, 2013) associated emotional valence with the rate of change of VFE. As in evaluative foraging, their framework introduces a meta-construction in that it applies to any active-inference process independently of the underlying generative model. Emotional valence is defined by the first and second derivatives of VFE, reflecting the dynamics of VFE. In contrast to evaluative foraging, the association between emotional valence and free energy was not extended to action, limiting its ability to explain or predict motivated behaviour. The reason is that this framework predated the development of EFE in active inference, which provides a formal mechanism to link affective states with action selection (Friston et al., 2015).

Emotional valence and adaptive behaviour have also been connected through active inference via the theory of *constructed emotion*, which posits that emotions emerge as holistic, context-dependent brain-body phenomena (Barrett, 2017). This theory highlights the role of predictive processes in constructing emotional experiences, with the brain dynamically modelling internal and external states to interpret sensory input and regulate the body. (Hesp et al., 2021) extended this framework by introducing *affective charge*, a computational perspective that incorporates meta-cognition, explicitly linking confidence in action selection to affective valence and subjective fitness through second-order beliefs (beliefs about beliefs). Affective charge can therefore be understood as a (signed) reward prediction error that increases the agent’s confidence in its action model if positive and decreases it if negative. However, it does not explain how behavioural decisions are made. Evaluative foraging, by contrast, emphasises the direct role of habit, wanting and liking in shaping behaviour. Affective charge can be integrated into evaluative foraging as second-order monitoring of reward maximisation, i.e. of free-energy minimisation.

The relationship between uncertainty reduction and positive hedonic value has also been explored as a mechanism for understanding art appreciation. These studies are framed within the theory of predictive coding, which serves as a key foundational component of the broader framework of active inference (Pezzulo et al., 2024). Predictive coding suggests that the brain seeks to make sense of sensory input by inferring the hidden states of the environment through Bayesian inference (Pezzulo et al., 2024). Applying this to art appreciation, (Van de Cruys, 2017) proposed that reducing prediction error by transitioning from higher uncertainty to lower uncertainty and positive rates of prediction error reduction elicit positive affective experiences.

Accordingly, the intense pleasure of “Aha” or *Erlebnis* moments arises from such a resolution of uncertainty. This perspective aligns with our proposal, where liking is proportional to the reduction in VFE. However, it does not explain why organisms engage in behaviours that can initially increase prediction error, as seen in exploratory or epistemic actions.

Overcoming this limitation, (Van de Cruys et al., 2024) recently proposed that engaging in behaviours that initially increase prediction error is explained by the epistemic component of curiosity, which reflects expected information gain and the expectation of resolving uncertainty. They formalise this as an *epistemic arc* spanning curiosity, epistemic action and resolution.

While their model draws on the process theory of active inference and acknowledges potential applications beyond art-related experiences (everyday sense-making processes), it remains largely conceptual and lacks a quantitative framework. Our approach addresses this gap by providing a quantitative method for characterising epistemic arcs, situating them within a continuum of experiences not restricted to specific domains or species. Evaluative foraging formalises the epistemic and pragmatic motivations and how hedonic evaluation drives them.

#### 5.1.2. Other models of hedonic value

Several approaches unrelated to active inference have proposed links between hedonic evaluation and motivated behaviour. (Skov, 2023) offered a conceptual model explaining how nervous systems assign hedonic value to sensory objects, leading to action. As in the current proposal, hedonic values are inherently adaptive, shaped by the organism’s survival needs and ever-changing environment. Liking and disliking trigger response patterns that enable organisms to recognise and react to situations that could aid or threaten their survival. Sensory valuation is not a simple reflection of an object’s intrinsic properties but depends on the organism’s idiosyncratic and current needs and evaluative context. While Skov’s model remains conceptual and does not accommodate data or make predictions, its conceptual similarities with our proposal and its detailed description of the neurobiology of hedonic evaluation make it a valuable resource for refining and enhancing descriptions reliant on computational theories such as active inference.

Evaluative foraging shares several features with computational models of hedonic value based on reinforcement learning, another theory of decision-making. In reinforcement learning, the objective is to determine the optimal policy (the one that maximises reward) through a value function following Bellman’s equation (Sutton & Barto, 2018), which includes immediate and future rewards. Focusing on hedonic value (referred to as *aesthetic* value), (Brielmann & Dayan, 2022) mapped two processes known to contribute to art appreciation to these two components of reinforcement learning: processing fluency (Reber et al., 2004) informs immediate rewards, while learning influences expected future rewards by facilitating future processing efficiency.

Drawing on this parallel, art experiences are processed in the brain similarly to how other rewarding experiences are handled in reinforcement learning, with hedonic value serving as a reward signal that guides future behaviour and preferences. While this proposal establishes a formal analogy with *wanting* and *liking* in hedonic evaluation, it does not account for intrinsic motivation for learning or pre-existing preferences driving behaviour. This limitation is inherent to all reinforcement learning models due to their lack of an explicit formulation of epistemic (uncertainty-reducing) and pragmatic (goal-directed) imperatives, at the core of active inference (K. J. Friston et al., 2017). In contrast with reinforcement learning, agents in active inference are endowed with a model of purposive behaviour: they act to fulfil their prior preferences. In addition, unlike value functions to be inferred, active inference naturally incorporates degrees of belief and uncertainty, central to understanding adaptive decision-making (Doya et al., 2007).

Therefore, active inference offers a compelling alternative to reinforcement learning for evaluating and generating motivated behaviour because it focuses on an agent’s beliefs, uncertainty and intrinsic motivations rather than externally defined rewards. As an active inference approach, evaluative foraging provides deeper insights into exploration, decision-making under uncertainty and adaptability in dynamic environments (Friston et al., 2016; Pezzulo et al., 2024; Schwartenbeck et al., 2019).

### 5.2. Mutual enrichment of hedonic evaluation and active inference

We have presented evaluative foraging as a mechanism through which reward maximisation drives behaviour in organisms with a reward system, with hedonic evaluation monitoring and motivating free-energy minimisation relative to specific, embodied affordances and conditions. We succinctly discuss the implications of merging hedonic evaluation and active inference in evaluative foraging, and how this integrated approach addresses the central questions raised in the introduction.

Through evaluative foraging, active inference makes several key contributions to hedonic evaluation. Traditionally, hedonic evaluation has included ways to quantify hedonic values but has lacked a formal mechanism for translating these values into decisions and actions. In our proposal, active inference fills this gap by providing hedonic evaluation with a direct computational link to motivated behaviour. By formalising hedonic evaluation within the active inference framework, researchers can systematically examine how the interplay of factors such as internal states and environmental cues leads to behavioural outcomes, and draw on current hypotheses about brain architecture in predictive processing and active inference to propose mechanisms for how hedonic values lead to action. Evaluative foraging unifies *wanting* and *liking*, the core components of hedonic evaluation, as complementary facets of a single objective: maximising reward, that is, minimising VFE. This approach embraces the *common currency* hypothesis in acknowledging a single evaluative mechanism regardless of reward and stimulus type, and transcends it by proposing a unified underlying mechanism for behaviour across complexity and taxa grounded in free-energy minimisation. In doing so, active inference frames hedonic evaluation within a first-principles account of behaviour, bridging neurobiological processes and computational goals.

Conversely, through our framework, hedonic evaluation offers significant insights with the potential to provide a useful perspective on active inference. The mechanisms underlying evaluation and reward are increasingly well defined in active inference. Reward maximisation is integrated through prior preferences that encode desired outcomes. These preferences guide policy selection by minimising EFE, which balances exploration and exploitation. Additionally, active inference formalises reward-related behaviour by treating prediction errors and learning rates as precision-weighted processes, akin to mechanisms in reinforcement learning but grounded in a unifying Bayesian framework. Dopamine has emerged as a key candidate for encoding precision-weighted prediction errors in active inference: phasic dopamine signals are thought to represent transient changes in precision, guiding learning and action by signalling reward prediction errors (Friston et al., 2012), while tonic dopamine signals (Bogacz, 2020), or the locus coeruleus–noradrenaline system (Sales et al., 2019), modulate the baseline confidence in policy selection. Despite these neurobiological formulations of active inference, much remains unknown about its precise neurobiological correlates. In particular, the neurobiological implementation of VFE minimisation is not so well formalised in the literature, and that of the correspondence between habits, EFE and VFE has not yet been clarified.

Thanks to its grounding in hedonic evaluation, evaluative foraging outlines possible directions to address these gaps by proposing for the first time a measurable proxy for free energy. As a first and paramount step, testing whether the activation of hedonic hotspots underlying liking corresponds to a reduction of VFE is crucial for understanding how behavioural choices are evaluated. Testing the hypotheses at the core of evaluative foraging as proposed in **Section 4.3** would provide valuable insights into the neurobiological correlates of active inference. Casting the constructs of hedonic evaluation – supported by its extensive neurobiological foundation (Berridge & Kringelbach, 2013, 2015) – within the active inference framework enables evaluative foraging to generate novel hypotheses and investigate the neurobiological mechanisms underlying free-energy minimisation.

In sum, evaluative foraging fosters a mutual enrichment of hedonic evaluation – the fundamental adaptive neurobiological process of assessing stimuli as beneficial or harmful – and active inference – a unified, principled theory of behaviour. This conceptual synthesis delivers two major contributions: First, evaluative foraging provides a principled explanation for motivated behaviour, establishing a causal link between hedonic values and action selection. Second, hedonic evaluation offers a biologically plausible measure for free-energy minimisation and a neuropsychological implementation of active inference. Together, evaluative foraging has the potential to explain how information is encoded in hedonic values, integrated and transformed into motivated behaviour, unifying computational and neurobiological mechanisms of behaviour into a single theoretical framework.

## 6. Conclusion

In this contribution, we propose a natural operationalisation of the main phases of hedonic evaluation and their corresponding value signals onto the main components of active inference. In our account, VFE corresponds to hedonic impact or core liking, quantifying sensory pleasure of ongoing experiences, while EFE captures the motivational drive or core wanting to pursue rewarding outcomes. This synthesis constitutes the foundation of our framework, evaluative foraging, which accounts for and predicts how hedonic evaluation generates and evaluates motivated behaviour.

Grounded in active inference, evaluative foraging provides a unifying framework for understanding preference-seeking behaviours through principles akin to those governing physical systems. Drawing on analogies from statistical physics, it conceptualises organisms as agents navigating probabilistic spaces to minimise VFE – similar to particles minimising a functional such as the Lagrangian (Friston, 2013). In this view, hedonic evaluation guides organisms along behavioural trajectories that minimise surprise, mirroring the principle of least action in physics.

This predictive framework relies on an embodied model of the world, shaped by an organism’s ecological niche, that generates expectations about and evaluates behavioural outcomes in the form of hedonic values. A key strength of evaluative foraging is its ability to unify diverse preference-seeking behaviours under a shared mechanistic framework, capturing a minimum common denominator in the motivational principles that drive behaviour. This extends the common currency hypothesis beyond static neural correlates of hedonic value to highlight the dynamic mechanisms underpinning hedonic choice in action planning. In doing so, our proposal bridges fundamental biological processes with higher-order cultural endeavours typically considered uniquely human, suggesting shared roots in adaptive mechanisms while accounting for the complexity of embodied, situated cognition and behaviour.

Evaluative foraging reinforces the idea that cognition is about engaging in purposive, adaptive interactions with the environment and its affordances (Pezzulo et al., 2024). This perspective represents a promising avenue for advancing artificial intelligence, where the move from traditional milestones in the field, such as the Turing test to the embodied Turing test (Zador et al., 2023), reflects this paradigm shift. We hope our proposal will inspire further investigation into the neurobiology of motivated behaviour, bridging foundational computational and biological mechanisms with complex cognitive functions and fostering interdisciplinary scientific progress.

## Acknowledgements

The authors are grateful to Fredrik Ullén for his critical review of a previous version of the manuscript. They also thank Marcus T. Pearce and Josep Marco-Pallarés for their valuable feedback, and Marcos Nadal, Martin Skov and Robert Zatorre for their encouraging reception of the idea.

## Footnotes

1 The processes and mechanisms discussed and proposed in this paper are not necessarily explicit or conscious, although they may be accompanied by explicit or conscious experience. Accordingly, we primarily refer to (implicit) core *wanting* (incentive salience) and *liking* (hedonic impact), as opposed to (explicit) conscious wanting (cognitive incentive) and liking (conscious pleasure) (Berridge & Dayan, 2021; Berridge & Kringelbach, 2015).

2 The concept of surprise in active inference originates in information theory and does not necessarily align with the concept of psychological surprise.

## References

Anderson, D. J., & Adolphs, R. (2014). A Framework for Studying Emotions across Species [Review]. Cell, 157(1), 187–200. 10.1016/j.cell.2014.03.003

Barnett, C. A., Bateson, M., & Rowe, C. (2007). State-dependent decision making: educated predators strategically trade off the costs and benefits of consuming aposematic prey [Article]. Behavioral Ecology, 18(4), 645–651. 10.1093/beheco/arm027

Barrett, L. F. (2017). The theory of constructed emotion: an active inference account of interoception and categorization [Article]. Social Cognitive and Affective Neuroscience, 12(1), 1–23. 10.1093/scan/nsw154

Barron, A. B., Sovik, E., & Cornish, J. L. (2010). The roles of dopamine and related compounds in reward-seeking behavior across animal phyla [Review]. Frontiers in Behavioral Neuroscience, 4, 9, Article 163. 10.3389/fnbeh.2010.00163

Bartra, O., McGuire, J. T., & Kable, J. W. (2013). The valuation system: A coordinate-based meta-analysis of BOLD fMRI experiments examining neural correlates of subjective value [Article]. Neuroimage, 76(1), 412–427. 10.1016/j.neuroimage.2013.02.063

Berridge, K. C. (1991). Modulation of taste affect by hunger, caloric satiety, and sensory-specific satiety in the rat [Article]. Appetite, 16(2), 103–120. 10.1016/0195-6663(91)90036-r

Berridge, K. C. (2004). Motivation concepts in behavioral neuroscience [Review]. Physiology and Behavior, 81(2), 179–209. 10.1016/j.physbeh.2004.02.004

Berridge, K. C., & Dayan, P. (2021). Liking [Editorial Material]. Current Biology, 31(24), R1555–R1555. 10.1016/j.cub.2021.09.069

Berridge, K. C., & Kringelbach, M. L. (2008). Affective neuroscience of pleasure: reward in humans and animals [Review]. Psychopharmacology, 199(3), 457–480. 10.1007/s00213-008-1099-6

Berridge, K. C., & Kringelbach, M. L. (2013). Neuroscience of affect: brain mechanisms of pleasure and displeasure [Review]. Current Opinion in Neurobiology, 23(3), 294–303. 10.1016/j.conb.2013.01.017

Berridge, K. C., & Kringelbach, M. L. (2015). Pleasure Systems in the Brain [Review]. Neuron, 86(3), 646–664. 10.1016/j.neuron.2015.02.018

Berridge, K. C., & Robinson, T. E. (1998). What is the role of dopamine in reward: hedonic impact, reward learning, or incentive salience? [Review]. Brain Research Reviews, 28(3), 309–369. 10.1016/s0165-0173(98)00019-8

Birch, L. L., & Marlin, D. W. (1982). I don’t like it; I never tried it: Effects of exposure on two-year-old children’s food preferences. Appetite, 3(4), 353–360. 10.1016/S0195-6663(82)80053-6

Bogacz, R. (2020). Dopamine role in learning and action inference [Article]. Elife, 9, 33, Article e53262. 10.7554/eLife.53262

Bornstein, R. F., & Dagostino, P. R. (1992). Stimulus-recognition and the mere exposure effect [Article]. Journal of Personality and Social Psychology, 63(4), 545–552. 10.1037//0022-3514.63.4.545

Brielmann, A. A., & Dayan, P. (2022). A Computational Model of Aesthetic Value [Article]. Psychological Review, 129(6), 1319–1337. 10.1037/rev0000337

Calhoun, A. J., Chalasani, S. H., & Sharpee, T. O. (2014). Maximally informative foraging by *Caenorhabditis elegans* [Article]. Elife, 3, 13, Article e04220. 10.7554/eLife.04220

Cannon, W. B. (1929). Organization for physiological homeostasis [Review]. Physiological Reviews, 9(3), 399–431. 10.1152/physrev.1929.9.3.399

Chiel, H. J., & Beer, R. D. (1997). The brain has a body: adaptive behavior emerges from interactions of nervous system, body and environment [Editorial Material]. Trends in Neurosciences, 20(12), 553–557. 10.1016/s0166-2236(97)01149-1

Cisek, P. (2022). Evolution of behavioural control from chordates to primates. Philosophical Transactions of the Royal Society B: Biological Sciences, 377(1844), 20200522. doi:10.1098/rstb.2020.0522

Clark, A. (2013). Whatever next? Predictive brains, situated agents, and the future of cognitive science [Article]. Behavioral and Brain Sciences, 36(3), 181–204. 10.1017/s0140525x12000477

Conant, R. C., & Ross Ashby, W. (1970). Every good regulator of a system must be a model of that system †. International Journal of Systems Science, 1(2), 89–97. 10.1080/00207727008920220

Costa, K. M., & Schoenbaum, G. (2022). Dopamine [Editorial Material]. Current Biology, 32(15), R817–R824. 10.1016/j.cub.2022.06.060

Darwin, C. (1872). The expression of the emotions in man and animals [doi:10.1037/10001-000]. John Murray. 10.1037/10001-000

Dayan, P. (2009). Goal-directed control and its antipodes [Article; Proceedings Paper]. Neural Networks, 22(3), 213–219. 10.1016/j.neunet.2009.03.004

Dayan, P. (2022). “Liking” as an early and editable draft of long-run affective value [Review]. PLoS Biology, 20(1), 15, Article e3001476. 10.1371/journal.pbio.3001476

Dolan, R. J., & Dayan, P. (2013). Goals and Habits in the Brain [Review]. Neuron, 80(2), 312–325. 10.1016/j.neuron.2013.09.007

Doya, K., Ishii, S., Pouget, A., & Rao, R. P. N. (2007). Bayesian brain: Probabilistic approaches to neural coding (K. Doya, S. Ishii, A. Pouget, & R. P. N. Rao, Eds.). MIT Press.

Drouin, A., Giguère, S., Déraspe, M., Marchand, M., Tyers, M., Loo, V. G., Bourgault, A. M., Laviolette, F., & Corbeil, J. (2016). Predictive computational phenotyping and biomarker discovery using reference-free genome comparisons [Article]. BMC Genomics, 17, 15, Article 754. 10.1186/s12864-016-2889-6

Duckworth, R. A. (2009). The role of behavior in evolution: a search for mechanism [Review]. Evolutionary Ecology, 23(4), 513–531. 10.1007/s10682-008-9252-6

FitzGerald, T. H. B., Dolan, R. J., & Friston, K. (2015). Dopamine, reward learning, and active inference [Article]. Frontiers in Computational Neuroscience, 9, 16, Article 136. 10.3389/fncom.2015.00136

Friedman, N. P., & Robbins, T. W. (2022). The role of prefrontal cortex in cognitive control and executive function [Review]. Neuropsychopharmacology, 47(1), 72–89. 10.1038/s41386-021-01132-0

Friston, K. (2013). Life as we know it [Article]. Journal of the Royal Society Interface, 10(86), 12, Article 20130475. 10.1098/rsif.2013.0475

Friston, K., FitzGerald, T., Rigoli, F., Schwartenbeck, P., O’Doherty, J., & Pezzulo, G. (2016). Active inference and learning [Review]. Neuroscience and Biobehavioral Reviews, 68, 862–879. 10.1016/j.neubiorev.2016.06.022

Friston, K., FitzGerald, T., Rigoli, F., Schwartenbeck, P., & Pezzulo, G. (2017). Active Inference: A Process Theory [Article]. Neural Computation, 29(1), 1–49. 10.1162/NECO_a_00912

Friston, K., Rigoli, F., Ognibene, D., Mathys, C., Fitzgerald, T., & Pezzulo, G. (2015). Active inference and epistemic value [Article]. Cognitive Neuroscience, 6(4), 187–214. 10.1080/17588928.2015.1020053

Friston, K. J. (2010). The free-energy principle: a unified brain theory? [Review]. Nature Reviews Neuroscience, 11(2), 127–138. 10.1038/nrn2787

Friston, K. J., Kilner, J., & Harrison, L. (2006). A free energy principle for the brain [Article]. Journal of Physiology-Paris, 100(1-3), 70–87. 10.1016/j.jphysparis.2006.10.001

Friston, K. J., Lin, M., Frith, C. D., Pezzulo, G., Hobson, J. A., & Ondobaka, S. (2017). Active Inference, Curiosity and Insight [Article]. Neural Computation, 29(10), 2633–2683. 10.1162/NECO_a_00999

Friston, K. J., Shiner, T., FitzGerald, T., Galea, J. M., Adams, R., Brown, H., Dolan, R. J., Moran, R., Stephan, K. E., & Bestmann, S. (2012). Dopamine, Affordance and Active Inference [Article]. PLoS Computational Biology, 8(1), 20, Article e1002327. 10.1371/journal.pcbi.1002327

Gartus, A., & Leder, H. (2014). The white cube of the museum versus the gray cube of the street: The role of context in aesthetic evaluations. Psychology of Aesthetics, Creativity, and the Arts, 8(3), 311–320. 10.1037/a0036847

Gibson, J. J. (1986). The Ecological Approach to Visual Perception. Lawrence Erlbaum Associates. https://books.google.fr/books?id=DrhCCWmJpWUC

Gintis, H. (2007). A framework for the unification of the behavioral sciences [Review]. Behavioral and Brain Sciences, 30(1), 1-+. 10.1017/s0140525x07000581

Gottlieb, J., & Oudeyer, P. Y. (2018). Towards a neuroscience of active sampling and curiosity [Review]. Nature Reviews Neuroscience, 19(12), 758–770. 10.1038/s41583-018-0078-0

Gregory, R. L. (1980). Perceptions as hypotheses [Article]. Philosophical Transactions of the Royal Society of London Series B-Biological Sciences, 290(1038), 181–197. 10.1098/rstb.1980.0090

Güntürkün, O., & Bugnyar, T. (2016). Cognition without Cortex [Review]. Trends in Cognitive Sciences, 20(4), 291–303. 10.1016/j.tics.2016.02.001

Hernández, M., Palomar-García, M. A., Nohales-Nieto, B., Olcina-Sempere, G., Villar-Rodríguez, E., Pastor, R., Avila, C., & Parcet, M. A. (2019). Separate Contribution of Striatum Volume and Pitch Discrimination to Individual Differences in Music Reward [Article]. Psychological Science, 30(9), 1352–1361, Article 0956797619859339. 10.1177/0956797619859339

Hesp, C., Smith, R., Parr, T., Allen, M., Friston, K. J., & Ramstead, M. J. D. (2021). Deeply Felt Affect: The Emergence of Valence in Deep Active Inference [Article]. Neural Computation, 33(2), 398–446. 10.1162/neco_a_01341

Hills, T. T., Todd, P. M., Lazer, D., Redish, A. D., Couzin, I. D., & Cognitive Search Res, G. (2015). Exploration versus exploitation in space, mind, and society. Trends in Cognitive Sciences, 19(1), 46–54. 10.1016/j.tics.2014.10.004

Jarvis, E. D., Gunturkun, O., Bruce, L., Csillag, A., Karten, H., Kuenzel, W., Medina, L., Paxinos, G., Perkel, D. J., Shimizu, T., Striedter, G., Wild, J. M., Ball, G. F., Dugas-Ford, J., Durand, S. E., Hough, G. E., Husband, S., Kubikova, L., Lee, D. W.,…Avian Brain Nomenclature, C. (2005). Avian brains and a new understanding of vertebrate brain evolution [Review]. Nature Reviews Neuroscience, 6(2), 151–159. 10.1038/nrn1606

Joffily, M., & Coricelli, G. (2013). Emotional Valence and the Free-Energy Principle [Article]. PLoS Computational Biology, 9(6), 14, Article e1003094. 10.1371/journal.pcbi.1003094

Kidd, C., & Hayden, B. Y. (2015). The Psychology and Neuroscience of Curiosity [Review]. Neuron, 88(3), 449–460. 10.1016/j.neuron.2015.09.010

Kleckner, I. R., Zhang, J. H., Touroutoglou, A., Chanes, L., Xia, C. J., Simmons, W. K., Quigley, K. S., Dickerson, B. C., & Barrett, L. F. (2017). Evidence for a large-scale brain system supporting allostasis and interoception in humans [Article]. Nature Human Behaviour, 1(5), 14, Article 0069. 10.1038/s41562-017-0069

Klink, P. C., van Wezel, R. J. A., & van Ee, R. (2012). United we sense, divided we fail: context-driven perception of ambiguous visual stimuli [Review]. Philosophical Transactions of the Royal Society B-Biological Sciences, 367(1591), 932–941. 10.1098/rstb.2011.0358

Koelsch, S., Vuust, P., & Friston, K. (2019). Predictive Processes and the Peculiar Case of Music [Review]. Trends in Cognitive Sciences, 23(1), 63–77. 10.1016/j.tics.2018.10.006

Kording, K. P., & Wolpert, D. M. (2006). Bayesian decision theory in sensorimotor control [Article; Proceedings Paper]. Trends in Cognitive Sciences, 10(7), 319–326. 10.1016/j.tics.2006.05.003

Kounios, J., & Beeman, M. (2014). The Cognitive Neuroscience of Insight. In S. T. Fiske (Ed.), Annual Review of Psychology, Vol 65 (Vol. 65, pp. 71–93). Annual Reviews. 10.1146/annurev-psych-010213-115154

Kringelbach, M. L. (2005). The human orbitofrontal cortex: Linking reward to hedonic experience [Review]. Nature Reviews Neuroscience, 6(9), 691–702. 10.1038/nrn1747

LeDoux, J. E. (2012). Evolution of human emotion: A view through fear. In M. A. Hofman & D. Falk (Eds.), Evolution of the Primate Brain: From Neuron to Behavior (Vol. 195, pp. 431–442). Elsevier Science Bv. 10.1016/b978-0-444-53860-4.00021-0

Levy, D. J., & Glimcher, P. W. (2012). The root of all value: a neural common currency for choice [Review]. Current Opinion in Neurobiology, 22(6), 1027–1038. 10.1016/j.conb.2012.06.001

Lipton, P. (2005). Testing hypotheses: Prediction and prejudice [Review]. Science, 307(5707), 219–221. 10.1126/science.1103024

Lopez-Persem, A., Domenech, P., & Pessiglione, M. (2016). How prior preferences determine decision-making frames and biases in the human brain [Article]. Elife, 5, 20, Article e20317. 10.7554/eLife.20317

Malekzadeh, P., & Plataniotis, K. N. (2024). Active Inference and Reinforcement Learning: A Unified Inference on Continuous State and Action Spaces Under Partial Observability [Article]. Neural Computation, 36(10), 2073–2135. 10.1162/neco_a_01698

Mappes, J., Marples, N., & Endler, J. A. (2005). The complex business of survival by aposematism [Review]. Trends in Ecology & Evolution, 20(11), 598–603. 10.1016/j.tree.2005.07.011

Marty, N., Dallaporta, M., & Thorens, B. (2007). Brain glucose sensing, counterregulation, and energy homeostasis [Review]. Physiology, 22, 241–251. 10.1152/physiol.00010.2007

Mesoudi, A., Whiten, A., & Laland, K. N. (2006). Towards a unified science of cultural evolution [Review]. Behavioral and Brain Sciences, 29(4), 329-+. 10.1017/s0140525x06009083

Montague, P. R., & King-Casas, B. (2007). Efficient statistics, common currencies and the problem of reward-harvesting [Review]. Trends in Cognitive Sciences, 11(12), 514–519. 10.1016/j.tics.2007.10.002

Nadal, M., & Skov, M. (2024). The sensory valuation account of aesthetic experience [Review; Early Access]. Nature Reviews Psychology, 15. 10.1038/s44159-024-00385-y

Nguyen, D., Naffziger, E. E., & Berridge, K. C. (2021). Positive affect: nature and brain bases of liking and wanting [Article]. Current Opinion in Behavioral Sciences, 39, 72–78. 10.1016/j.cobeha.2021.02.013

O’Connell, L. A., & Hofmann, H. A. (2011). The Vertebrate mesolimbic reward system and social behavior network: A comparative synthesis [Review]. Journal of Comparative Neurology, 519(18), 3599–3639. 10.1002/cne.22735

Panksepp, J. (1998). The periconscious substrates of consciousness: Affective states and the evolutionary origins of the self. Journal of Consciousness Studies, 5(5-6), 566–582.

Parr, T., Pezzulo, G., & Friston, K. J. (2022). Active inference: the free energy principle in mind, brain, and behavior. MIT Press.

Patzelt, E. H., Hartley, C. A., & Gershman, S. J. (2018). Computational Phenotyping: Using Models to Understand Individual Differences in Personality, Development, and Mental Illness. Personal Neurosci, 1, e18. 10.1017/pen.2018.14

Penacchio, O., Halpin, C. G., Cuthill, I. C., Lovell, P. G., Wheelwright, M., Skelhorn, J., Rowe, C., & Harris, J. M. (2024). A computational neuroscience framework for quantifying warning signals [Article]. Methods in Ecology and Evolution, 15(1), 103–116. 10.1111/2041-210x.14268

Penacchio, O., Hämäläinen, L., Rojas, B., Summers, K., Yeager, J., Sherratt, T. N., & Exnerová, A. (2025). Cognitive ecology of surprise in predator-prey interactions [Review; Early Access]. Functional Ecology, 17. 10.1111/1365-2435.14750

Pereira, C. S., Teixeira, J., Figueiredo, P., Xavier, J., Castro, S. L., & Brattico, E. (2011). Music and Emotions in the Brain: Familiarity Matters [Article]. PloS One, 6(11), 9, Article e27241. 10.1371/journal.pone.0027241

Perry, C. J., & Barron, A. B. (2013). Neural Mechanisms of Reward in Insects. In M. R. Berenbaum (Ed.), Annual Review of Entomology, Vol 58 (Vol. 58, pp. 543–562). Annual Reviews. 10.1146/annurev-ento-120811-153631

Pessiglione, M., & Lebreton, M. (2015). From the Reward Circuit to the Valuation System: How the Brain Motivates Behavior. In G. H. E. Gendolla, M. Tops, & S. L. Koole (Eds.), Handbook of Biobehavioral Approaches to Self-Regulation (pp. 157–173). Springer New York. 10.1007/978-1-4939-1236-0_11

Pezzulo, G., & Cisek, P. (2016). Navigating the Affordance Landscape: Feedback Control as a Process Model of Behavior and Cognition [Review]. Trends in Cognitive Sciences, 20(6), 414–424. 10.1016/j.tics.2016.03.013

Pezzulo, G., Parr, T., & Friston, K. (2024). Active inference as a theory of sentient behavior [Article]. Biological Psychology, 186, 9, Article 108741. 10.1016/j.biopsycho.2023.108741

Pezzulo, G., Rigoli, F., & Friston, K. (2015). Active Inference, homeostatic regulation and adaptive behavioural control [Review]. Progress in Neurobiology, 134, 17–35. 10.1016/j.pneurobio.2015.09.001

Pitman, R. L., & Durban, J. W. (2012). Cooperative hunting behavior, prey selectivity and prey handling by pack ice killer whales (Orcinus orca), type B, in Antarctic Peninsula waters [Article]. Marine Mammal Science, 28(1), 16–36. 10.1111/j.1748-7692.2010.00453.x

Ramsay, D. S., & Woods, S. C. (2014). Clarifying the Roles of Homeostasis and Allostasis in Physiological Regulation [Article]. Psychological Review, 121(2), 225–247. 10.1037/a0035942

Rangel, A., Camerer, C., & Montague, P. R. (2008). A framework for studying the neurobiology of value-based decision making [Review]. Nature Reviews Neuroscience, 9(7), 545–556. 10.1038/nrn2357

Reber, R., Schwarz, N., & Winkielman, P. (2004). Processing fluency and aesthetic pleasure: Is beauty in the perceiver’s processing experience? [Review]. Personality and Social Psychology Review, 8(4), 364–382. 10.1207/s15327957pspr0804_3

Reber, R., Winkielman, P., & Schwarz, N. (1998). Effects of perceptual fluency on affective judgments [Article]. Psychological Science, 9(1), 45–48. 10.1111/1467-9280.00008

Roh, E., Song, D. K., & Kim, M. S. (2016). Emerging role of the brain in the homeostatic regulation of energy and glucose metabolism [Review]. Experimental and Molecular Medicine, 48, 12, Article e216. 10.1038/emm.2016.4

Rosenthal, G. G. (2017). Mate choice: The evolution of sexual decision making from microbes to humans. Princeton University Press.

Ross, S., & Hansen, N. C. (2016). Dissociating Prediction Failure: Considerations from Music Perception [Editorial Material]. Journal of Neuroscience, 36(11), 3103–3105. 10.1523/jneurosci.0053-16.2016

Roth, W. M., & Jornet, A. (2013). Situated cognition [Article]. Wiley Interdisciplinary Reviews-Cognitive Science, 4(5), 463–478. 10.1002/wcs.1242

Sales, A. C., Friston, K. J., Jones, M. W., Pickering, A. E., & Moran, R. J. (2019). Locus Coeruleus tracking of prediction errors optimises cognitive flexibility: An Active Inference model [Article]. PLoS Computational Biology, 15(1), 24, Article e1006267. 10.1371/journal.pcbi.1006267

Schlaug, G. (2015). Musicians and music making as a model for the study of brain plasticity. In E. Altenmuller, S. Finger, & F. Boller (Eds.), Music, Neurology, and Neuroscience: Evolution, the Musical Brain, Medical Conditions, and Therapies (Vol. 217, pp. 37–55). Elsevier Science Bv. 10.1016/bs.pbr.2014.11.020

Schultz, W. (1998). Predictive reward signal of dopamine neurons [Review]. Journal of Neurophysiology, 80(1), 1–27. 10.1152/jn.1998.80.1.1

Schultz, W. (2015). Neuronal reward and decision signals: from theories to data [Review]. Physiological Reviews, 95(3), 853–951. 10.1152/physrev.00023.2014

Schurr, R., Reznik, D., Hillman, H., Bhui, R., & Gershman, S. J. (2024). Dynamic computational phenotyping of human cognition [Article]. Nature Human Behaviour, 8(5), 18. 10.1038/s41562-024-01814-x

Schwartenbeck, P., & Friston, K. (2016). Computational Phenotyping in Psychiatry: A Worked Example [Article]. Eneuro, 3(4), 18, Article 0049-16.2016. 10.1523/eneuro.0049-16.2016

Schwartenbeck, P., Passecker, J., Hauser, T. U., FitzGerald, T. H. B., Kronbichler, M., & Friston, K. J. (2019). Computational mechanisms of curiosity and goal-directed exploration [Article]. Elife, 8, 45, Article e41703. 10.7554/eLife.41703

Schwarz, N. (2007). Attitude construction: Evaluation in context [Article]. Social Cognition, 25(5), 638–656. 10.1521/soco.2007.25.5.638

Seger, C. A., & Spiering, B. J. (2011). A Critical Review of Habit Learning and the Basal Ganglia [Hypothesis and Theory]. Frontiers in Systems Neuroscience, volume 5 - 2011. 10.3389/fnsys.2011.00066

Sescousse, G., Caldú, X., Segura, B., & Dreher, J. C. (2013). Processing of primary and secondary rewards: A quantitative meta-analysis and review of human functional neuroimaging studies [Review]. Neuroscience and Biobehavioral Reviews, 37(4), 681–696. 10.1016/j.neubiorev.2013.02.002

Shapiro, L. (2011). Embodied Cognition. Routledge. <GO to ISI>://WOS:000399995800009

Shettleworth, S. J. (2012). Do Animals Have Insight, and What Is Insight Anyway? [Article]. Canadian Journal of Experimental Psychology-Revue Canadienne De Psychologie Experimentale, 66(4), 217–226. 10.1037/a0030674

Skelhorn, J., & Rowe, C. (2006). Prey palatability influences predator learning and memory [Article]. Animal Behaviour, 71, 1111–1118. 10.1016/j.anbehav.2005.08.011

Skov, M. (2023). Sensory liking: How nervous systems assign hedonic value to sensory objects. In M. S. M. Nadal (Ed.), The Routledge international handbook of neuroaesthetics (pp. 31–62). Routledge.

Smith, R., Badcock, P., & Friston, K. J. (2021). Recent advances in the application of predictive coding and active inference models within clinical neuroscience [Article]. Psychiatry and Clinical Neurosciences, 75(1), 3–13. 10.1111/pcn.13138

Smith, R., Friston, K. J., & Whyte, C. J. (2022). A step-by-step tutorial on active inference and its application to empirical data [Article]. Journal of Mathematical Psychology, 107, 60, Article 102632. 10.1016/j.jmp.2021.102632

Snell-Rood, E. C. (2013). An overview of the evolutionary causes and consequences of behavioural plasticity [Article]. Animal Behaviour, 85(5), 1004–1011. 10.1016/j.anbehav.2012.12.031

Spaan, M. T. J. (2012). Partially Observable Markov Decision Processes. In M. Wiering & M. van Otterlo (Eds.), Reinforcement Learning: State-of-the-Art (pp. 387–414). Springer Berlin Heidelberg. 10.1007/978-3-642-27645-3_12

Sterling, P. (2012). Allostasis: A model of predictive regulation [Article]. Physiology and Behavior, 106(1), 5–15. 10.1016/j.physbeh.2011.06.004

Sutton, R. S., & Barto, A. G. (2018). Reinforcement Learning: An Introduction, 2nd Edition. Mit Press. <GO to ISI>://WOS:000481873900019

Theriault, J. E., Katsumi, Y., Reimann, H. M., Zhang, J., Deming, P., Dickerson, B. C., Quigley, K. S., & Barrett, L. F. (2025). It’s not the thought that counts: Allostasis at the core of brain function. Neuron. 10.1016/j.neuron.2025.09.028

Thorndike, E. L. (1911). Animal intelligence: Experimental studies [doi:10.5962/bhl.title.55072]. Macmillan Press. 10.5962/bhl.title.55072

Topolinski, S., & Reber, R. (2010). Gaining Insight Into the “Aha” Experience [Article]. Current Directions in Psychological Science, 19(6), 402–405. 10.1177/0963721410388803

Turk-Browne, N. B. (2012). Statistical Learning in Perception. In N. M. Seel (Ed.), Encyclopedia of the Sciences of Learning (pp. 3182–3185). Springer US. 10.1007/978-1-4419-1428-6_1707

Unoki, S., Matsumoto, Y., & Mizunami, M. (2006). Roles of octopaminergic and dopaminergic neurons in mediating reward and punishment signals in insect visual learning [Article]. European Journal of Neuroscience, 24(7), 2031–2038. 10.1111/j.1460-9568.2006.05099.x

Van de Cruys, S. (2017). Affective Value in the Predictive Mind. In T. K. Metzinger & W. Wiese (Eds.), Philosophy and Predictive Processing. MIND Group. 10.15502/9783958573253

Van de Cruys, S., Frascaroli, J., & Friston, K. (2024). Order and change in art: towards an active inference account of aesthetic experience [Review]. Philosophical Transactions of the Royal Society B-Biological Sciences, 379(1895), 15, Article 20220411. 10.1098/rstb.2022.0411

von Helmholtz, H. (1866). Concerning the perceptions in general (Vol. 3). Dover.

Wiener, N. (1948). Cybernetics; or control and communication in the animal and the machine. John Wiley.

Winkielman, P., & Berridge, K. (2003). Irrational wanting and subrational liking: How rudimentary motivational and affective processes shape preferences and choices [Review]. Political Psychology, 24(4), 657–680. 10.1046/j.1467-9221.2003.00346.x

Xiang, T., Ray, D., Lohrenz, T., Dayan, P., & Montague, P. R. (2012). Computational Phenotyping of Two-Person Interactions Reveals Differential Neural Response to Depth-of-Thought [Article]. PLoS Computational Biology, 8(12), 9, Article e1002841. 10.1371/journal.pcbi.1002841

Yamamoto, K., & Vernier, P. (2011). The evolution of dopamine systems in chordates [Review]. Frontiers in Neuroanatomy, 5, 21, Article 21. 10.3389/fnana.2011.00021

Zador, A., Escola, S., Richards, B., Ölveczky, B., Bengio, Y., Boahen, K., Botvinick, M., Chklovskii, D., Churchland, A., Clopath, C., DiCarlo, J., Ganguli, S., Hawkins, J., Körding, K., Koulakov, A., LeCun, Y., Lillicrap, T., Marblestone, A., Olshausen, B.,…Tsao, D. (2023). Catalyzing next-generation Artificial Intelligence through NeuroAI [Article]. Nature Communications, 14(1), 7, Article 1597. 10.1038/s41467-023-37180-x

Zajonc, R. B. (1968). Attitudinal effects of mere exposure [Article]. Journal of Personality and Social Psychology, 9(2P2), 1-&. 10.1037/h0025848

Zatorre, R. J., Chen, J. L., & Penhune, V. B. (2007). When the brain plays music: auditory-motor interactions in music perception and production [Review]. Nature Reviews Neuroscience, 8(7), 547–558. 10.1038/nrn2152

